# Lineage-determining transcription factors constrain cohesin to drive multi-enhancer oncogene regulation

**DOI:** 10.1101/2025.10.10.681717

**Authors:** Yeqiao Zhou, Atishay Jay, Noah Burget, Tobias Friedrich, Sora Yoon, Jessica Alsing, Guy Nir, Rudolf Grosschedl, Golnaz Vahedi, Robert B. Faryabi

## Abstract

Multiple enhancers, often located across vast genomic distances, regulate key genes. However, how chromatin topology organization at individual alleles enables cell-type-restricted multi-enhancer gene regulation remains unclear. Using acute protein degradation and time-course population-average chromatin conformation capture in lymphoma, we found that the B-cell-lineage-determining transcription factor EBF1 preferentially positions multiple enhancers at loci containing sparsely distributed genes essential for B-cell identity and oncogenesis. Our time-resolved sub-diffraction optical chromatin architecture tracing of >100,000 alleles in individual lymphoma cells further revealed diverse topological conformations facilitating multi-enhancer interactions. Mechanistically, we found that positioning of enhancers at allelic topological centers is required for their interactions with target promoters, with EBF1 serving as a barrier to the loop-extruding cohesin on enhancers. These findings, which we demonstrate their generalizability to the T-cell-lineage-determining transcription factor TCF1 in T-cell leukemia, suggest that lineage-determining transcription factors radially position enhancers and promoters to enable multi-enhancer regulation of key oncogenes.

## INTRODUCTION

Precise spatiotemporal control of gene expression relies on the activities of *cis*-regulatory elements, including enhancers and promoters^1-6^. Earlier studies indicate that multiple enhancers, sometimes thousands of base pairs away on the linear genome, can be in the spatial proximity of one target gene promoter, contributing to its transcription in normal and malignant cells^7-11^. However, the mechanisms by which the physical folding of individual chromatin fiber facilitates the formation of multi-enhancer-promoter interactions remain unclear. Additionally, although the activation of specific enhancers has been attributed to lineage-determining transcription factors^12-20^, the role of these factors in organizing chromatin fiber folding to enable multi-enhancer gene regulation remains elusive.

Interaction networks among multiple enhancers and promoters were first implicated by chromatin conformation capture methods with progressively higher resolutions. These techniques, such as Hi-C, HiChIP and Micro-C, are powerful sequencing-based tools for profiling population-average genome-wide chromatin organization features^21-26^. However, they obscure the behavior of individual alleles within cells and are further limited to detecting pairwise interactions in fragmented chromatin. By contrast, Oligopaint DNA fluorescence *in situ* hybridization (FISH), a diffraction-limited imaging method, has been used to demonstrate spatial positioning of a few regulatory elements in individual cancer cells^27-29^. Yet, this approach is restricted by the availability of fluorescent channels, and similar to sequencing-based assays lacks information about folding of entire loci. Thus, it fails to reveal the relationship between positioning of enhancers and promoters and their surrounding chromatin. To overcome these limitations and elucidate how chromatin fiber folding at individual alleles enables cancer type-restricted multi-enhancer gene regulation, we implemented Optical Reconstruction of Chromatin Architecture (ORCA), a sequential DNA FISH method, to examine the precise folding of whole genomic regions in individual cancer cells at sub-diffraction resolution^30,31^.

The application of ORCA at the *Sox9* and *Pitx1* loci has revealed the diverse individual structures underlying changes in chromatin folding during development^32,33^. However, these studies primarily focused on topologically associating domains (TADs) and the regulation of TAD structures by ubiquitously expressed architectural proteins such as the ring-shaped cohesin complex and the insulator protein CCCTC-binding factor (CTCF). Unlike TAD boundaries, which are generally cell-type invariant, enhancer activities and positioning are mostly cell-lineage and disease specific^2,3,11,25,27,34^. Notably, recent protein degradation studies have shown that loss of cohesin and CTCF profoundly alters TAD structures with limited effect on enhancer-promoter interactions and transcription in cell populations^35,36^. These observations sparked interest in investigating the roles of other transcription factors in lineage- and disease-specific regulation of chromatin folding.

While lineage-determining transcription factors are known to activate lineage-specific enhancers that drive gene expression in normal and cancer cells^37-39^, their role in the spatial positioning of enhancers is less understood. Recent studies of normal cells demonstrated that the cooperativity of T-cell-lineage-determining transcription factor TCF1 and CTCF impacts TAD boundaries during T cell development^40^. Stem cell reprogramming studies further showed that the pluripotent factor KLF4 contributes to chromatin reorganization and rewiring of enhancer-promoter interactions^41^. Extending similar studies to cancer, we and others showed that NOTCH1, a key signaling-dependent oncogene and T-cell-lineage-instructing transcription factor, contributes to reorganization of enhancer-promoter interactions in Notch-addicted cancers^27,42^. These population-level studies provide evidence that non-architectural transcription factors contribute to chromatin organization in general and enhancer positioning in particular. However, how lineage-determining transcription factors organize chromatin folding at an entire locus to instruct multi-enhancer gene regulation in both normal and malignant cells remains unknown.

In this study, we optically traced the chromatin architectures of four loci in mantel cell lymphoma (MCL) and T acute lymphoblastic leukemia (T-ALL) to examine how organization of chromatin topology of individual alleles enables cancer type-restricted multi-enhancer gene regulation. By time-resolved tracing of over 100,000 chromatin fibers, we discovered that irrespective of their linear genomic locations, the placement of distal enhancers and promoters to allelic spatial centers is required for their interactions. We demonstrated that B and T cell lineage-determining transcription factors, EBF1^38^ and TCF1^43^, bind cancer-type-restricted enhancers and act as barriers to cohesin traffic, thereby restricting loop extrusion and promoting the centralization of interacting enhancers and promoters. Our single-allele studies further revealed that gene activation alone is not sufficient, but rather transcription factor-dependent distal enhancer activation is required for central positioning of enhancers and promoters. These findings provide fundamental insights into how multi-enhancer-promoter interactions form in chromatin and suggest a general mechanism by which lineage-determining transcription factors alter spatial positioning of regulatory elements to control gene expression, contributing to oncogenesis.

## RESULTS

### EBF1 binds distal enhancers and promotes MCL survival

To elucidate the role of lineage-determining transcription factors in cancer-type-restricted multi-enhancer gene regulation, we begin with examining the dependency of B cell lymphomas to *EBF1*, a transcription factor that is required for B-lineage commitment of hematopoietic precursors^44-46^. Our analysis of published data from 1,237 cancer cell lines^47^ showed that EBF1 was highly expressed and selectively essential for growth of various Non-Hodgkin B cell lymphomas, including mantle cell lymphoma (MCL) (**Fig. S1A)**. CRISPR-Cas9-mediated genomic deletion of EBF1 (hereafter referred to as EBF1 knockout) in MCL models supported these observations and further showed that loss of highly expressed EBF1 (**Fig. S1B)** induced apoptosis and cell death in MCL (**Figs. S1C and S1D**).

EBF1 is a key regulator of chromatin structure and gene expression during normal B-cell differentiation^38,48^. Observing EBF1 essentiality for MCL survival prompted us to examine the chromatin activity of EBF1-bound genomic elements in MCL Granta519. A total of 25,227 high-confidence EBF1-bound elements, identified using two independent EBF1 antibodies, were enriched for enhancer histone marks H3K27ac and H3K4me1, and were devoid of the Polycomb histone mark H3K27me3 (**Fig. 1A and Table S1**). Additionally, we observed that 75% of these EBF1-bound elements were more than 10 Kb from active gene promoters (**Fig. S1E**). Taken together, these data suggest that EBF1 predominantly binds distal enhancers in MCL.

**Figure 1:**
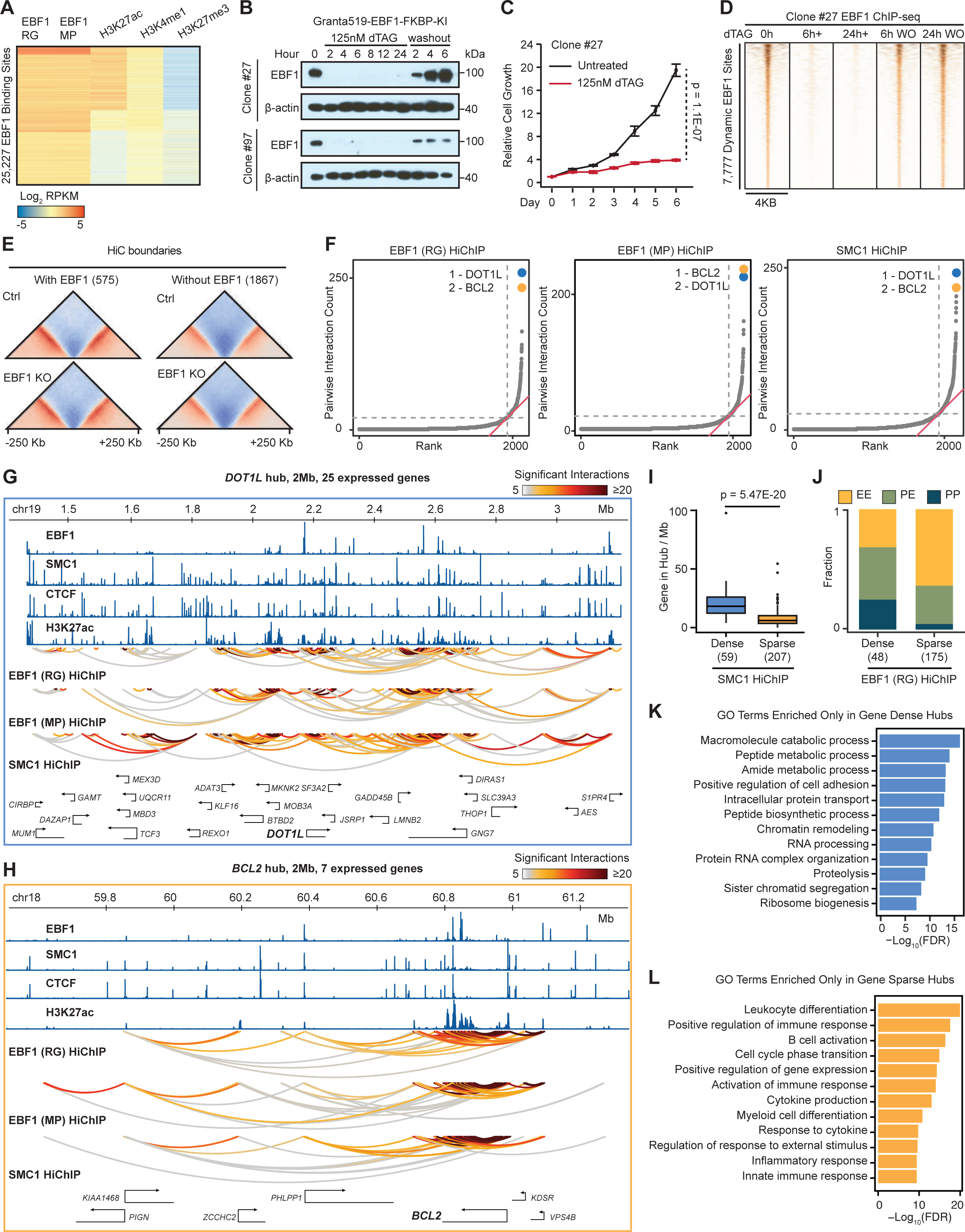
EBF1 binds distal enhancers in gene-dense and gene-sparse hubs. A: Heatmap displaying enrichment of active histone marks at 25,227 reproducible EBF1 binding sites determined by two ChIP-grade antibodies (RG and MP). EBF1 binding sites are ordered by hierarchical clustering of normalized EBF1, H3K27ac, H3K4me1 and H3K27me3 ChIP-seq signals. B: Western blotting of EBF1 in EBF1-FKBP-KI Granta519 clones 27 and 97 showing rapid and reproducible removal and restoration of EBF1 protein levels after dTAG^V^-1 (referred to as dTAG) treatment and washout, respectively. 125 nM dTAG treatment and washout are examined for 0 to 24 hours. β-actin is loading control. C: EBF1 loss impedes growth of Granta519 clone 27. Relative cell growth (CellTiter-glo) of clone 27 with or without 125 nM dTAG treatment for 6 days. Data represent mean ± SEM of n=5 biological replicates. P-value: t-test. D: Heatmap of EBF1 occupancy showing 7,777 bona fide EBF1-bound elements determined with Log2(fold change) < -1 in EBF1-removed (6h+dTAG and 24h+dTAG versus 0h) and Log2(fold change) > 1 in EBF1-restored (6h WO and 24h WO versus 24h+dTAG) Granta519 clone 27. Each column is centered on EBF1-bound elements +/- 2 Kb flanking sequences with 50 bp resolution. WO: washout. E: Pileup plots showing TAD boundaries are invariant in Granta519-Cas9 cells with (top) and without (bottom) EBF1 expression. Left: centered around 575 EBF1-bound TAD boundaries. Right: 1,867 EBF1-unbound boundaries. F: Population-average enhancer-promoter hubs defined by EBF1 and SMC1 HiChIP in Granta519. Groups of interacting enhancers and/or promoters are plotted in an ascending order of their total pairwise interactions from EBF1 HiChIP using RG antibody (left) and MP antibody (middle) and SMC1 HiChIP (right). Population-average enhancer-promoter hubs are defined as the groups of enhancers and promoters with total pairwise interactions above the elbow of total interaction distribution. The top two hubs of each HiChIP are marked and named with their representative genes. RG: a polyclonal anti-EBF1 antibody recognizing an N-terminal EBF1 peptide. MP: Millipore anti-EBF1 antibody. G, H: Gene-dense *DOT1L* and gene-sparse *BCL2* hubs display distinct distributions of enhancers and promoters and loops across the same linear genomic spans. ChIP-seq tracks showing enrichment of EBF1, SMC1, CTCF, and H3K27ac at proximal and distal enhancers of *DOT1L* (G) and *BCL2* (H), which are the top two highest interacting hubs defined in (F). HiChIP arcs displaying normalized significant interactions among enhancers and promoters. Bottom track indicating positions of expressed Ensembl genes in Granta519. I: Boxplot showing higher number of genes per megabase in gene-dense compared to gene-sparse hubs defined by SMC1 HiChIP. p-values: Wilcoxon rank sum test. Number of hubs in each group is listed in parenthesis. J: Barplots depicting the fraction of interactions among enhancers (E) and promoters (P) in gene-dense and gene-sparse hubs defined by EBF1 HiChIP, RG antibody. Number of hubs in each group is listed in parentheses. K, L: Gene Ontology terms specifically enriched in gene-dense (K) or gene-sparse (L) hubs. The top 500 expressed genes in each class of hubs are used as input.

### dTAG-induced EBF1 degradation and restoration rapidly alter EBF1 chromatin binding in MCL

Observing that EBF1 binds MCL distal enhancers led us to examine its immediate effects on enhancer activity and positioning. To this end, we used the degradation tag (dTAG) system^49^ and generated ∼100 independent MCL Granta519 clones, in which FKBP^F36V^ degradation tag was knocked into the C-terminus of both endogenous alleles of *EBF1* (hereafter referred to as EBF1- FKBP-KI MCL) (**Fig. S1F**). Among our independent EBF1-FKBP-KI lines, clones 97 and 27, which were used for all subsequent studies, exhibited the highest transcriptomic similarity to parental Granta519 cells (**Fig. S1G, Table S2**).

Temporal immunoblotting analysis of clones 97 and 27 with a polyclonal anti-EBF1 antibody (RG)^38^, which recognizes an N-terminal EBF1 peptide, showed that our degron strategy resulted in the acute removal and full restoration of EBF1 as early as 2 hours after a dTAG ligand (dTAG^V^- 1) addition and washout, respectively (**Fig. 1B**). As expected, long-term acute EBF1 degradation phenocopied EBF1 genetic deletion and blunted MCL proliferation (**Fig. 1C**). More importantly, this analysis revealed that, in contrast to long-term removal, 24-hour acute EBF1 degradation had a marginal impact on EBF1-FKBP-KI MCL proliferation (**Fig. 1C**).

Given that EBF1 chromatin binding precedes chromatin opening by 18 hours in early B-cell differentiation^50^, we used chromatin immunoprecipitation sequencing (ChIP-seq) to examine the dynamics of EBF1 chromatin occupancy in EBF1-FKBP-KI MCL. Notably, this analysis identified 7,777 bona fide EBF1 binding sites where chromatin-bound EBF1 was efficiently removed and restored following 6-hour dTAG ligand addition and washout, respectively (**Fig. 1D and Table S1**). Together, these data demonstrate that the EBF1 degron is a tractable and robust strategy for resolving the direct effects of EBF1 on MCL chromatin structure and topology on a timescale of hours without affecting cell proliferation. Hence, we conducted all subsequent experiments, unless otherwise noted, at short time points (6 and 24 hours).

### MCL chromatin accessibility is insensitive to rapid EBF1 changes

Chromatin structure and topology are both crucial for proper gene expression control^8,11,51-53^. Given that EBF1 induces lineage-specific chromatin accessibility in lymphoid progenitors during B-cell specification^38,54^, we first examined the immediate impact of EBF1 on MCL chromatin structure. Time-resolved ATAC-seq studies in EBF1-FKBP-KI MCL revealed a substantial loss of chromatin accessibility 48 hours after EBF1 removal (**Figs. S1H and S1I; Table S3**) when proliferation was significantly affected (**Fig. 1C**). By contrast, EBF1 removal and restoration did not markedly alter chromatin accessibility at 6 and 24 hours following dTAG ligand addition and washout (**Figs. S1H and S1I; Table S3**). Together, these data suggest that the immediate impact of EBF1 on MCL chromatin structure is marginal.

### MCL topologically associating domains and compartments are insensitive to EBF1

Compartments and topologically associating domains (TADs) are megabase-scale, population-level features of chromatin architecture, influencing enhancer activity and specificity by contributing to their spatial positioning^1,55,56^. Given that EBF1 predominantly binds MCL distal enhancers, where accessibility levels are largely invariant to rapid EBF1 changes (**Figs. 1A, S1E, S1H, and S1I; Table S3**), we next examined the impact of EBF1 on TAD boundary elements and compartments (**Table S4**). Analysis with *in situ* Hi-C (600 million sequencing reads per library), an assay that is frequently used to delineate TADs but is limited in detecting high-resolution regulatory loops, showed that EBF1 knockout did not impact the insulation potential of EBF1- occupied TAD boundaries (**Figs. 1E and S1B**). To further examine the impact of EBF1 on chromatin architecture, we next compared Hi-C-defined euchromatic A and heterochromatic B compartments in EBF1 wild-type and knockout MCL cells (**Fig. S1J**). Similar to TAD structures, loss of EBF1 for 3 days did not markedly alter the compartmentalization of the MCL genome. Taken together, these data suggest that compartments and TAD boundaries are invariant to rapid EBF1 changes across MCL populations.

### EBF1-involved interactions participate in MCL population-average enhancer-promoter hubs

Observing that EBF1 loss does not restructure compartments and TADs, we postulated that EBF1 instructs ensemble enhancer-promoter hubs, defined as a network of interacting enhancers and promoters across the population, but not necessarily simultaneously within individual cells. To evaluate this hypothesis, we examined interactions involving EBF1 and the extruder cohesin complex using HiChIP, an assay that enables interrogation of interactions attributed to a specific protein. HiChIP with a previously validated antibody against the cohesin complex component SMC1a^23,27^ (830 million sequencing reads) identified 26,363 significant high-resolution (∼5 Kb) SMC1-involved loops between locus pairs. Given there was no precedent for EBF1 HiChIP, we used two independent antibodies (RG and MP: Millipore) with ∼1 billion sequenced reads per experiment to identify significant high-resolution (∼5 Kb) EBF1-involved loops.

To investigate a potential interplay between EBF1 and ensemble enhancer-promoter hubs, we first compared the H3K27ac load as well as EBF1 and YY1 chromatin occupancy levels at enhancer and promoter elements interacting via cohesin-involved loops (**Fig. S1K**). As expected, H3K27ac was equally present at enhancers and promoters that interacted with other enhancers or promoters. YY1, which is known to preferentially bind promoters^57^, was more abundant at promoter-promoter interactions (**Fig. S1K**). By contrast, EBF1 preferentially bound to enhancers that interact with other enhancers (**Fig. S1K**), suggesting a potential role for EBF1 in mediating enhancer-enhancer and/or enhancer-promoter interactions.

Intrigued by this observation, we leveraged SMC1 and EBF1 HiChIP to further examine the relationship between EBF1 and MCL ensemble enhancer-promoter hubs (**Fig. 1F and Table S4**). These studies revealed that interacting enhancers and promoters identified via cohesin or EBF1 HiChIP were asymmetrically distributed, with only a small subset of these elements forming ensemble enhancer-promoter hubs characterized by a substantial number of pairwise interactions (**Fig. 1F and Table S4**). Analysis of cohesin-involved loops in MCL showed that while 1,849 of 2,115 loci containing enhancers and/or promoters had fewer than 23 pairwise interactions, 12.6% of loci (266 loci) participated in ensemble enhancer-promoter hubs with interactivity levels reaching up to 250 pairwise interactions (**Fig. 1F**). Similarly, EBF1-involved interactions formed 186 (MP) and 223 (RG) hubs, with 171 shared between the two libraries and 166 also marked as hubs by SMC1 HiChIP (**Fig. S1L**), as exemplified by two of the most interacting hubs containing the histone methyltransferase *DOT1L*^58^ and the anti-apoptotic factor *BCL2*^59^ (**Fig. 1F**). Taken in conjunction, these findings suggest a high degree of concordance between highly interacting enhancers and promotes groups that coalesce through EBF1- and cohesin-involved loops.

### EBF1-bound gene-dense and gene-sparse hubs encompass distinct classes of genes

To better understand potential regulatory environments formed by EBF1-bound ensemble hubs in MCL, we closely inspected *DOT1L* and *BCL2* regions (**Figs. 1G and 1H**), the two hubs with the highest number of EBF1- and cohesin-involved pairwise interactions (**Fig. 1F**). While both *DOT1L*- and *BCL2*-containing hubs consisted of interactions among highly acetylated EBF1- and SMC1-bound elements and spanned comparable linear genomic regions, the gene-dense *DOT1L* hub harbored 3.5 times more expressed genes than the gene-sparse *BCL2* hub. Extending this analysis genome-wide revealed that fewer than 25% of the EBF1-bound hubs were located in gene-dense regions, with more than 15 active genes per hub (**Fig. S1M**). Similar to the *DOT1L* hub, gene-dense hubs harbored disproportionately higher densities of hub-positioned genes, with nearly 4 times more genes per megabase compared to gene-sparse hubs (median 16 vs 4 genes) (**Figs. 1I and S1N**).

In line with these observations, a comparison of these two classes of ensemble enhancer-promoter hubs revealed that more than 50% of pairwise interactions in gene-sparse hubs occurred between enhancers (**Fig. 1J**). By contrast, over 70% of interactions in gene-dense hubs were either between an enhancer and a promoter or between two promoters (**Fig. 1J**), suggesting that gene-dense and gene-sparse hubs may be architecturally distinct. To further explore this conjecture, we tested whether these two classes of enhancer-promoter hubs were associated with the regulation of different molecular functions in MCL. Notably, genes within gene-dense hubs function in general cellular processes such as chromatin remodeling and RNA processing (**Fig. 1K**), including *GAPDH*, *ACLY*, *BRD4* and *EEF1A2*. In contrast, the genes within gene-sparse hubs encoded proteins involved in B-cell-related pathways (**Fig. 1L**), as exemplified by *PAX5*, *BCL2L11*, *IKZF1* and *EBF1* itself^60-62^, further supporting the potential structure-function differences between gene-dense and gene-sparse hubs.

### EBF1 preferentially regulates genes within MCL population-average gene-sparse hubs

Observing the differential positioning of lineage-related and housekeeping genes in EBF1-bound gene-sparse and gene-dense hubs led us to next examine whether EBF1 differentially organized these two classes of enhancer-promoter hubs. Consistent with the higher number of enhancers and promoters in gene-dense hubs, we observed more EBF1-bound regulatory elements in gene-dense than in gene-sparse hubs (**Figs. 2A and S2A**). However, EBF1 binding at enhancers and promoters was significantly more dynamic in gene-sparse hubs 6 and 24 hours after dTAG ligand addition (**Fig. 2B**) and washout (**Fig. 2C**). In line with these observations, changes in EBF1 significantly and reproducibly altered the expression of genes located in gene-sparse compared with gene-dense hubs (**Figs. 2D and S2B**), as exemplified by rapid changes in the proto-oncogene *MYC* **(Figs. 2E and 2F)** and the PD-L1-encoding gene *CD274* (**Figs. S2C-E**). Together, these population-average data hint that EBF1 preferentially regulates genes positioned in gene-sparse hubs.

**Figure 2:**
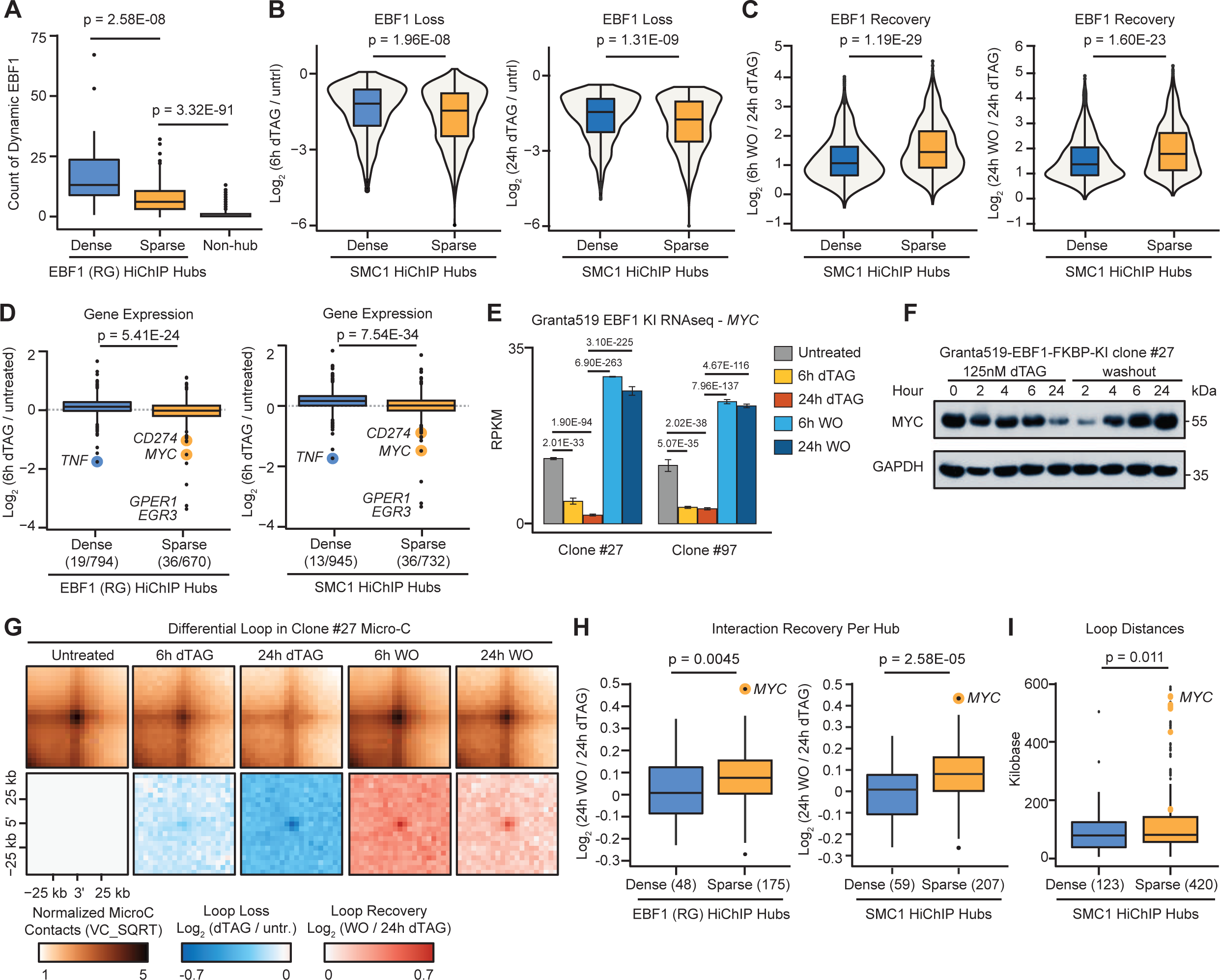
EBF1 preferentially activates target genes in gene-sparse hubs by promoting interactions with distal enhancers. A: Boxplots comparing number of EBF1 binding sites in gene-dense, gene-sparse and non-hub regions, as determined with EBF1 HiChIP. P-values: Wilcoxon rank sum test. B: Box and violin plots showing significantly higher loss of EBF1 binding in gene-sparse compared to gene-dense hubs 6 (left) and 24 (right) hours after dTAG treatment. P-values: Wilcoxon rank sum test. C: Box and violin plots showing significantly higher recovery of EBF1 binding in gene-sparse compared to gene-dense hubs 6 (left) and 24 (right) hours after dTAG washout. P-values: Wilcoxon rank sum test. D: Boxplots showing more significant down-regulation of genes in gene-sparse compared to gene-dense hubs defined by EBF1 (left) and SMC1 (right) HiChIP 6 hours after dTAG treatment. Note that there are fewer expressed genes in gene-sparse hubs. Numbers of differential / expressed genes in each group are listed in parentheses. E: Barplots of normalized RNAseq reads showing significant *MYC* down- and up-regulation 6 and 24 hours after dTAG treatment and washout, respectively, in Granta519 EBF1-FKBP-KI clones 27 and 97. Data represent mean ± S.D. of 3 replicates per condition. FDR from DESeq2. F: Time-course western blotting showing MYC protein levels in Granta519 EBF1-FKBP-KI clone 27 after dTAG treatment and washout. GAPDH is loading control. G: Pileup plots of normalized time-course Micro-C contact maps of clone 27 showing interactions among enhancers and promoters in EBF1 HiChIP hubs decrease upon dTAG treatment and recover after dTAG washout. Each heatmap showing +/- 25 Kb flanking the center with 5 Kb resolution. H: Boxplots showing significantly stronger recovery of Micro-C interactions among enhancers and promoters 24 hours after dTAG washout in gene-sparse compared to gene-dense hubs as defined by EBF1 (left) and SMC1 (right) HiChIP. Each dot represents the average Log2(fold change) of 24h washout over 24h dTAG of interactions in each hub. Number of hubs in each group is listed in parentheses. I: Boxplots showing longer genomic distances of interactions among enhancers and promoters in gene-sparse compared to gene-dense hubs. Yellow dots represent interactions of the *MYC* hub. Number of hubs in each group is listed in parentheses.

### EBF1 preferentially instructs MCL population-average gene-sparse hubs

Encouraged by this observation, we directly examined whether EBF1 differentially impacts the genome architecture of gene-sparse and gene-dense hubs. Given Hi-C’s limited resolution and HiChIP’s dependency on EBF1 or SMC1 protein binding, we performed Micro-C, an MNase-based chromatin conformation assay that enables genome-wide and unbiased mapping of population-average enhancer and/or promoter interactions^21,22^. Time-resolved Micro-C showed that EBF1 removal and restoration gradually decreased and increased the frequency of looping between enhancers and/or promoters in MCL hubs, respectively (**Fig. 2G**). Although the number of enhancer and/or promoter loops with at least two-fold change in interaction frequency was not markedly different between these two classes of hubs (**Fig. S2F**), the magnitude of interaction frequency changes upon EBF1 removal and restoration was greater in gene-sparse hubs (**Figs. 2H and S2G**), where we observed longer loops among enhancers and/or promoters (**Fig. 2I**). Taken together with our EBF1-dependent gene expression analysis (**Figs. 2D-F, S2B-E**), our genome-wide chromatin conformation capture data suggest a preferential role for EBF1 in controlling structure-function of population-average gene-sparse hubs in MCL.

### Optical chromatin tracing of individual *MYC* alleles in MCL reveals pronounced EBF1- instructed topological changes

Our population-average chromatin conformation capture data showed that rapid and acute changes in EBF1 have the most pronounced effect on pairwise interactions at the *MYC* gene-sparse hub **(Figs. 2H and S2G**), which comprises some of the longest loops in the EBF1-FKBP- KI MCL genome (**Fig. 2I**). Given that the *MYC* hub exemplifies the differential impact of EBF1 on gene-sparse and gene-dense hubs, we closely examined its population-average looping using EBF1 and SMC1 HiChIP. We observed frequent pairwise interactions between the *MYC* promoter and the two bona fide *MYC* super-enhancers E1 and E2 located 525 and 433 Kb 5’ of the promoter, respectively^63^ **(Fig. 3A)**. However, these data cannot determine whether the *MYC* hub is an emergent property of population-averaged pairwise interactions across different cells or if EBF1-instructed multi-way enhancer-promoter interactions occur simultaneously in individual MCL cells. To answer this question and precisely examine the impact of EBF1 on the topology of the *MYC* gene-sparse hub at the level of individual chromatin fibers, we conducted time-course ORCA, a multiplexed FISH methodology that relies on sequential hybridization and imaging steps for super-resolution measurement of distances between all segments along a chromatin trace^31^ (**Fig. 3B**).

**Figure 3:**
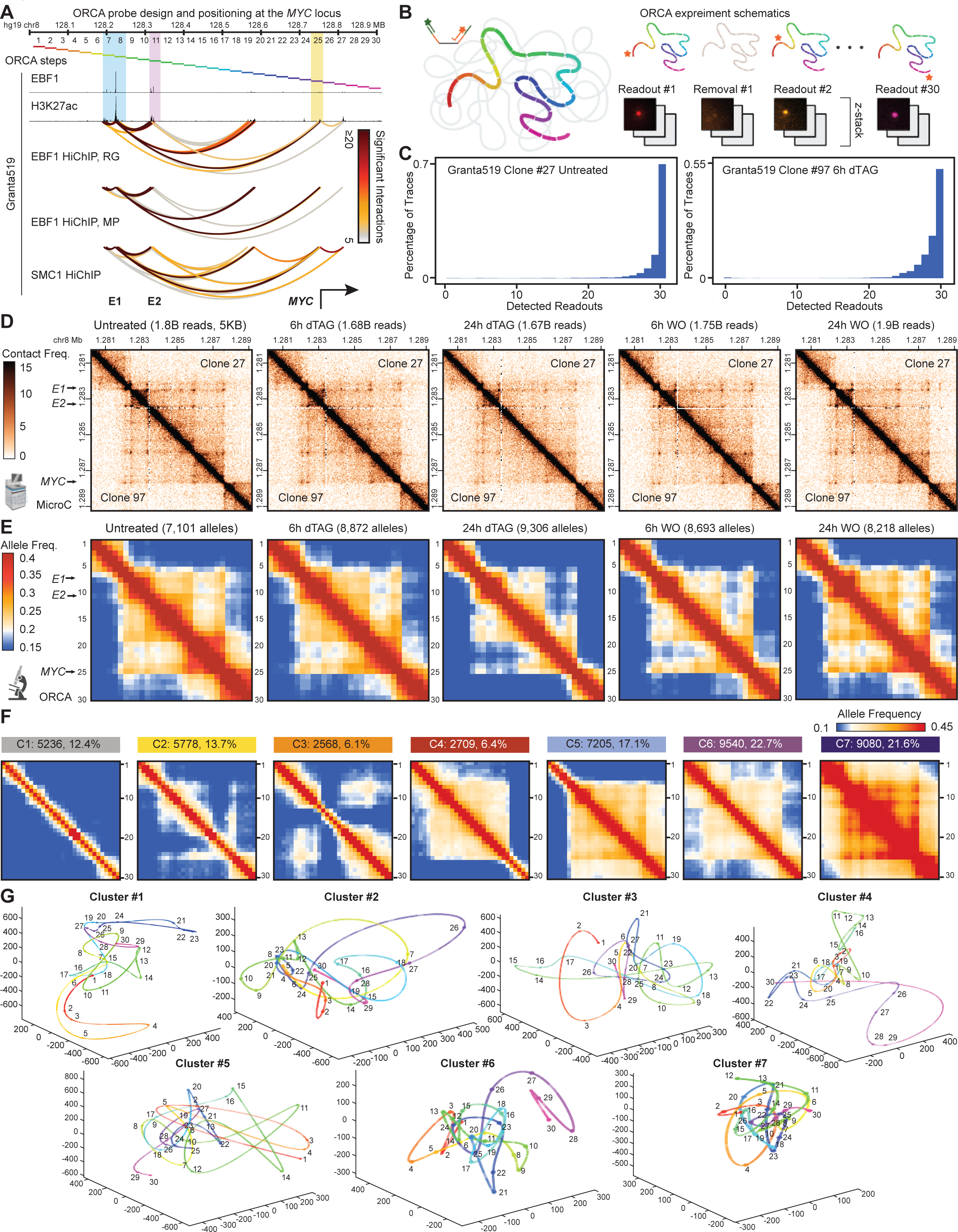
Single-allele optical tracing of the *MYC* locus reveals EBF1-instructed topological changes. A: ChIP-seq tracks showing EBF1 and H3K27ac at the two distal super-enhancers E1 and E2 located 525 and 433 Kb 5’ of the *MYC* promoter in MCL. SMC1 and EBF1 HiChIP arcs displaying normalized significant interactions among promoters and enhancers in Granta519. Bottom track indicates E1, E2 and *MYC* gene location. Top track indicates the 30 steps of ORCA experiments. B: Schematic of ORCA experiment procedures at the *MYC* locus in Granta519. C: Representative barplots showing the majority of traces have all 30 segment readouts detected in the ORCA experiments of clone 27 (left) and clone 97 (right) in untreated (left), 6h dTAG treatment (right), 24h dTAG treatment, 6h dTAG washout and 24h dTAG washout conditions (**Table S6**). Only traces with <= 5 missing barcodes are used in remaining analyses. D: Time-course Micro-C interaction frequency maps showing gradual and reproducible (top: clone 27, bottom: clone 97) reduction and recovery of interactions at the *MYC* locus after dTAG treatment and washout. Resolution: 5 Kb. E: Time-course ORCA allele frequency maps showing gradual reduction and recovery of interactions at the *MYC* locus after dTAG treatment and washout. Each condition combines the alleles from clones 27 and 97 and uses the average distance between two consecutive segments as the cutoff for determining interaction. Alleles are not imputed, and missing values are excluded from the calculations. F: Allele frequency maps of *MYC* locus traces for each of the 7 common topological conformations. Clustering and calculation of frequencies are performed on pairwise distance matrices of imputed alleles from all conditions in the two clones. G: Example reconstructed chromatin traces for each of the 7 common topological conformations of the *MYC* locus. Numbers on the traces indicate the 30 ORCA steps.

To examine the topology of individual *MYC* alleles while balancing genome coverage and throughput, we designed 5,891 primary oligonucleotide probes and employed our modified version of the ORCA protocol to tile a 900-Kb region containing the *MYC* promoter (**Table S5**), its two super-enhancers, as well as two strong nearby TAD boundaries and their flanking sequences at 30-Kb resolution (**Fig. 3A**). At each step of sequential imaging, we efficiently labeled, imaged, and subsequently removed the uniquely barcoded probes of a given 30-Kb segment (**Fig. S3A**). This process was repeated until all 30 segments were precisely localized (**Fig. S3B**). The registration of high-accuracy fiducial signals, which continuously marked alleles throughout sequential imaging (**Fig. S3A**), enabled super-resolution measurement of the three-dimensional positions of all 30 segments in >60% of chromatin traces (**Fig. 3C and Table S6**), demonstrating markedly higher efficiency than earlier studies^32,33^.

Using our optimized ORCA protocol, we visualized ∼43,000 chromatin traces from untreated, 6-hour and 24-hour dTAG treated, as well as 6-hour and 24-hour dTAG washout EBF1-FKBP-KI MCL clones 27 and 97. Given there was no gap between adjacent segments, we used the average distance between two consecutive segments instead of an arbitrary number as the cutoff for determining interactions (i.e. loops) between any two segments along a chromatin trace. Given the marked reproducibility of population-average pairwise interaction frequency maps (**Table S6**), we merged data from thousands of individual chromatin traces from EBF1-FKBP-KI MCL clones 27 and 97. In all five conditions, the population-average ORCA and Micro-C-detected pairwise interaction frequencies were highly correlated (**Figs. S3C and S3D**). Furthermore, the *MYC* TAD / sub-TAD boundaries and enhancer-promoter loop dots were clearly observed in ORCA’s average distance and interaction frequency maps, aligning with those identified by Micro-C (**Figs. 3D, 3E, S3C and S3D**).

Additionally, ORCA average distance and interaction frequency maps revealed more pronounced changes in the topology of the *MYC* locus. The centromeric and telomeric sub-TADs progressively separated and interacted less frequently 6 and 24 hours after EBF1 removal (**Figs. 3E and S3C**). Notably, a stripe-like feature emanating from the ORCA probe sets overlapping the *MYC* promoter, super-enhancer E1, and super-enhancer E2 gradually diminished 6 and 24 hours after EBF1 removal (**Figs. 3E and S3C**). Furthermore, time-course ORCA experiments showed that EBF1 removal steadily reduced interactions across the telomeric TAD boundary of the locus, separating the *MYC* promoter from its 3’ flanking sequences (**Figs. 3E and S3C**). More importantly, ORCA average distance and interaction frequency maps showed that EBF1 restoration gradually reinstated interactions between centromeric and telomeric sub-TADs, the stripe from the *MYC* promoter to its distal super-enhancers, as well as cross-telomeric TAD boundary interactions. Together, these data confirm the high quality of our ORCA data and demonstrated that, even at the population level, ORCA provides additional insights into the impact of EBF1 on *MYC* locus topology in MCL.

### EBF1 alters the frequency of *MYC* common topological conformations in MCL

Given the clear EBF1-dependency of population-average *MYC* locus interaction frequency maps (**Figs. 3E and S3C**), we next asked whether EBF1 instructs a unique chromatin conformation of the *MYC* locus, or changes the frequency of common topological conformations available to *MYC*. To begin differentiating between these two hypotheses, we first leveraged the single-allele nature of ORCA data and clustered high-dimensional chromatin traces from all EBF1-FKBP-KI MCL cells. This analysis identified seven distinct common conformations, each with more than 2,000 *MYC* chromatin traces (**Figs. 3F and S3E**). These common conformations displayed varying levels of compaction, with the most open and compact traces grouped together in conformations C1 and C7, respectively. No interaction between the *MYC* promoter and its two super-enhancers was observed in traces with conformation C1. By contrast, the *MYC* promoter and super-enhancers E1 and E2 frequently interacted through loops and stripes in the absence of TAD-like features in the traces grouped in conformations C2 and C3, respectively. The traces with the conformation C4 exhibited marked interactions between E1 and E2 and their centromeric flanking sequences, as shown by a high frequency of interactions across the centromeric TAD boundary. Conversely, traces with conformation C5 showed a high degree of *MYC* interaction with its telomeric flanking sequences, reflected in a high frequency of interactions across the telomeric TAD boundary and a low frequency across the centromeric TAD boundary. Notably, only slightly more than 20% of traces assumed conformation C6, exhibiting features reminiscent of population-average structures of EBF1-expressing MCL cells, including prominent *MYC*-to-E1 and *MYC*-to-E2 interactions and stripe. Together, these data suggest that the *MYC* promoter samples from a set of common topological conformations despite the heterogeneity in *MYC* locus folding.

We next performed a similar analysis in our time-course ORCA experiments to examine the impact of EBF1 on common conformations of the *MYC* locus. Notably, while the set of possible common *MYC* chromatin conformations remained invariant (**Figs. S4A-C**), EBF1 removal and restoration rapidly altered the prevalence of traces with certain chromatin conformations (**Fig. S3E**). For instance, frequency of traces with conformation C2, where the *MYC* promoter and super-enhancers E1 and E2 interacting through a stripe-like feature, decreased from 14.3% to less than 8% of the alleles 24 hours after EBF1 removal. Notably, the percentage of traces with conformation C5 gradually decreased from ∼22% to 12.6% after EBF1 removal and subsequently recovered upon EBF1 restoration. On the other hand, the frequency of traces with conformation C4, which is topologically in contrast with C5, increased to 18.4% 24 hours after EBF1 removal and returned to the level of untreated cells (10%) after EBF1 restoration. Taken together, our results support a model in which EBF1 does not alter the constellation of topologies available to *MYC* but rather increases the likelihood of conformations that are permissive to interactions between *MYC* and its distal enhancers (**Fig. S3E**), resulting in substantial EBF1-dependent differences in population-average interaction frequency maps (**Fig. 3E**).

Examination of heterogeneity of pairwise interaction frequency maps revealed common conformations of the *MYC* locus and their EBF1-dependency in MCL cells. To complement this analysis, we visually inspected topological features of individual alleles using Optical Looping Interactive Viewing Engine (OLIVE), a portal that we developed for interactive querying and exploration of chromatin traces^64^. Notably, our visual inspection revealed that despite being at opposite ends of the locus on the linear genome (**Fig. 3A**), the *MYC* promoter and its two super-enhancers tended to locate at the spatial centers of chromatin traces with common conformations permissive to promoter-enhancer interaction (**Fig. 3G**). Intrigued by this observation, we quantified the geometric center of each chromatin trace and measured how far each segment along the trace was from the center (**Fig. S3F**). In line with our visual observations, this quantitative analysis showed that traces with different common conformations had distinct local radial patterns. Notably, the *MYC* promoter was, in general, closer to the geometric center in traces with conformations that had a higher tendency of enhancer-promoter interactions (**Figs. S3F and 3G**). Collectively, our data suggest a potential link between relative radial positioning of the *MYC* promoter within the locus and its ability to interact with its distal enhancers.

### EBF1 instructs *MYC* multi-enhancer interactions in individual MCL cells

Our analysis revealed that EBF1 alters the frequency of common topological conformations assumed by the *MYC* locus, potentially in relation to the promoter’s local radial positioning. Intrigued by this observation, we sought to examine how EBF1 impacts heterogeneity of E1 and E2 positioning relative to the *MYC* promoter. We observed a lack of coordination in *MYC* promoter-enhancer interactions between alleles of the same nucleus under all conditions (**Fig. 4A**). Thus, we first determined the frequencies of alleles with *MYC*-to-E1 interactions from our time-course ORCA experiments (**Fig. 4B**). As an internal control, we also quantified interaction frequencies between distance-matched non-enhancer/promoter pairs across the locus using a sliding window (**Fig. 4B**). This analysis revealed that *MYC* and E1 interacted in 37% of alleles, a frequency that steadily and significantly decreased after EBF1 removal. Notably, 24 hours after EBF1 degradation, the frequency of alleles with *MYC*-to-E1 interactions was comparable to that of distance-matched internal controls. Conversely, EBF1 restoration rapidly and significantly increased the frequency of alleles with *MYC*-to-E1 interactions within 6 hours, reaching levels comparable to those observed in dTAG-untreated cells (**Fig. 4B**).

**Figure 4:**
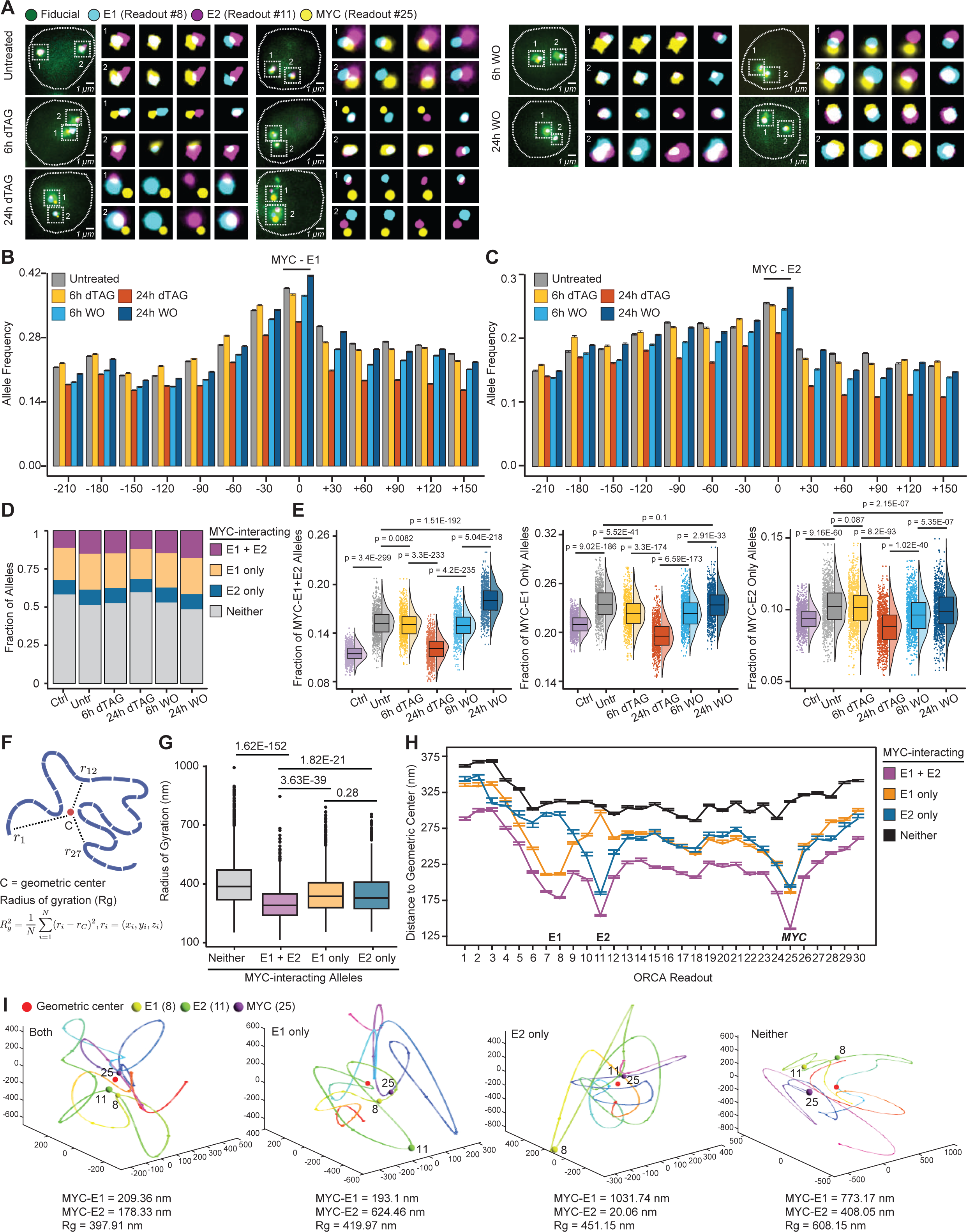
The *MYC* promoter and enhancers interact at the allelic geometric centers in MCL. A: Distances between the *MYC* promoter, E1 and E2 significantly increase and decrease after dTAG treatment and washout, respectively. Two representative nuclei per condition are shown with fiducial and magnified images for *MYC*-E1-E2, *MYC*-E1, *MYC*-E2 and E1-E2. To distinguish the steps imaged with Alexa647 sequentially, E1, E2 and MYC are pseudo-colored as cyan, magenta and yellow, and fiducial using Cy3 is pseudo-colored as green. B: Barplots of allele frequencies with E1 (step 7 or 8) and *MYC* (step 25) interactions compared with the distance-matched pairs using a sliding window (see Methods). Bootstrapping is performed per condition and error bars show 95% confidence intervals. C: Same as B for E2 (step 11) and *MYC* (step 25). D: Barplots showing fraction of alleles with *MYC*-E1-E2, only *MYC*-E1, only *MYC*-E2 or no *MYC* promoter-enhancer interaction (Neither) per condition. Control regions are defined in Methods. E: Box and violin plots showing decrease and recovery of allele frequency with *MYC*-E1-E2, only *MYC*-E1 and only *MYC*-E2 interactions after dTAG treatment and washout. Each point represents the allele frequencies calculated from 500 randomly sample alleles with a total of 1000 rounds of random sampling per condition. P-values: Wilcoxon rank sum test. F: Definition of geometric center and radius of gyration. G: Boxplots showing distribution of radius of gyration of alleles with *MYC*-E1-E2, only *MYC*-E1, only *MYC*-E2 or no *MYC* promoter-enhancer interaction (Neither) in untreated MCL. P-values: Wilcoxon rank sum test. H: Distances of each segment to the geometric centers of chromatin traces with *MYC*-E1-E2, only *MYC*-E1, only *MYC*-E2 or no *MYC* promoter-enhancer interactions (Neither). The median values of stratified alleles are calculated and plotted for each ORCA segment in untreated cells. Error bars show 95% confidence interval from bootstrapping. I: Example reconstructed chromatin traces for alleles with *MYC*-E1-E2 three-way interaction, *MYC*-E1 interaction, *MYC*-E2 interaction and no interaction emphasizing the central positioning of interacting elements. Rg: radius of gyration.

We next turned to E2 and examined its positioning relative to the *MYC* promoter at individual alleles. The allele frequency of pairwise *MYC*-to-E2 interactions was 24% in untreated cells, which decreased to a level comparable to that of distance-matched internal controls 24 hours after EBF1 removal (**Fig. 4C**). Examination of interactions after dTAG washout revealed that EBF1 rapidly restored *MYC*-to-E2 interactions to an allele frequency comparable to that of untreated cells within 6 hours (**Fig. 4C**). Collectively, these single-allele resolution studies complemented our population-average Micro-C observations and further demonstrate that EBF1 binding to *MYC* super-enhancers decreases *MYC* promoter-enhancer proximity in individual MCL chromatin fibers.

In light of these single-allele resolution observations, we next examined whether multi-enhancer-promoter interactions are present in individual chromatin fibers. To this aim, we quantified the frequency of alleles with only pairwise *MYC*-to-E1, *MYC*-to-E2, or three-way *MYC*-E1-E2 interactions. This analysis revealed that *MYC*-E1-E2 simultaneous interactions occur in 12% of alleles at any given time (**Fig. 4D**). Although relatively rare, *MYC*-E1-E2 interactions were twice as frequent as expected based on interactions among distance-matched non-enhancer/promoter element triplets (**Fig. 4D**). Bootstrapping analysis further substantiated the statistical significance of this observation (**Fig. 4E**). Taken in conjunction, these data demonstrate that single-allele enhancer-promoter hubs are formed in individual MCL cells, potentially by co-occurring dynamic interactions among enhancers and promoters.

Observing multi-enhancer interactions at individual *MYC* alleles led us to investigate the role of EBF1 in their formation. To this end, we compared the frequency of alleles with *MYC*-E1-E2 three-way interactions in untreated, 6- and 24-hours dTAG treated, and 6- and 24-hours dTAG washout cells. This analysis revealed that the loss of EBF1 gradually decreased three-way interaction frequencies between the *MYC* promoter and its two distal super-enhancers across individual alleles (**Figs. 4D and 4E**). 24 hours after EBF1 removal, the frequency of alleles with *MYC* multi-enhancer interactions significantly reduced to a level comparable to that in distance-matched internal controls (**Figs. 4D and 4E**). Conversely, we observed that EBF1 recovery rapidly and significantly increased the frequency of multi-enhancer-promoter interactions within 6 hours, reaching levels comparable to those observed in dTAG-untreated cells.

Notably, we observed differences in both the magnitude and dynamics of recovery of three-way compared to pairwise interactions (**Fig. 4E**). EBF1 restoration increased allele frequencies for both pairwise and three-way interactions; however, the establishment of *MYC*-E1-E2 three-way interactions was markedly more rapid and prevalent. Six hours after EBF1 restoration, the *MYC*- E1-E2 allele frequency was comparable to that in untreated cells and continued to increase over the following 18 hours. By contrast, the frequencies of alleles with only *MYC*-to-E1 or *MYC*-to-E2 pairwise interactions were lower at the 6-hour washout compared to untreated cells and only recovered to comparable levels 24 hours after EBF1 restoration. In summary, our findings demonstrate that in addition to facilitating pairwise enhancer-promoter interactions in individual MCL cells, EBF1 contributes to the maintenance and establishment of multi-enhancer interactions, an EBF1-preferred topological conformation that enables rapid *MYC* induction.

### The *MYC* promoter and enhancers interact at the allelic spatial centers in MCL

Our single-allele data have advanced our earlier population-level studies, elucidating that the lineage-determining transcription factor EBF1 instructs multi-enhancer interactions to control *MYC* expression in MCL. However, how the folding of chromatin fibers at individual alleles enables multi-enhancer gene regulation remains unclear. Additionally, it is unknown how lineage-determining transcription factors impact the interplay between these two levels of spatial genome organization.

Intrigued by our observation that the local radial positioning of the *MYC* promoter coincides with the locus conformation permissiveness to enhancer-promoter interactions (**Fig. S3F**), we hypothesized that the centering of the *MYC* promoter and enhancers influences their multiway interactions. To test this, we stratified chromatin traces based on the status of *MYC* promoter interactions with E1 and E2. The radius of gyration (**Fig. 4F**), a measure of individual chromatin fiber compaction, showed that traces containing *MYC* multi-enhancer interactions were markedly more compact than traces with only *MYC*-to-E1, only *MYC*-to-E2, or no *MYC* promoter-enhancer interactions (**Figs. 4G, 4I, and S4D**). More importantly, examination of traces with multi-enhancer interactions revealed that *MYC*, E1 and E2 - despite being located at the 3’ or 5’ ends of the locus on the linear genome - were markedly placed toward the geometric center relative to the rest of the region (**Figs. 4H, 4I, and S4D**).

In contrast, segments of chromatin traces with no enhancer-promoter interactions exhibited uniformly larger distances from the locus center (**Figs. 4H, 4I, and S4D**). Notably, E1 was markedly displaced from the locus’s spatial center in traces with only *MYC*-to-E2 interactions, while E2 was similarly separated from the center in traces with only *MYC*-to-E1 interactions (**Figs. 4H, 4I, and S4D**). Importantly, these observations remained consistent despite changes in allele frequency for *MYC*-to-E1 and/or *MYC*-to-E2 interactions following EBF1 degradation and restoration (**Figs. S4E-H**).

Taken together, these findings, along with the observation that EBF1 increases the frequency of alleles with *MYC* promoter-enhancer interactions (**Figs. 4B-E**), suggest that EBF1 contributes to the local radial positioning of enhancers to facilitate multi-enhancer-promoter interactions in individual MCL cells.

### EBF1 selectively impacts cohesin distribution on MCL chromatin

It has been shown that architectural proteins, including the cohesin complex, CTCF, and YY1, facilitate interactions between enhancers and promoters^65-67^. To gain insights into the underlying mechanisms of EBF1-mediated positioning of promoters and enhancers at the geometrical center of the *MYC* locus, we first investigated the impact of EBF1 on the chromatin occupancy of these proteins in EBF1-FKBP-KI MCL cells after EBF1 removal and restoration. Changes in CTCF and YY1 chromatin occupancy did not correlate with EBF1 binding loss (**Fig. S5A**). By contrast, SMC1 chromatin occupancy and EBF1 binding alterations correlated after dTAG treatment and washout (**Fig. 5A**). Consistent with the observation that EBF1 has a greater impact on looping in gene-sparse than gene-dense hubs (**Figs. 2G-I and S2G**), we observed markedly greater changes in SMC1 loading in gene-sparse than gene-dense hubs 6 and 24 hours after EBF1 removal and restoration (**Figs. 5B and S5B**). Given that SMC1 levels were invariant to EBF1 (**Fig. 5C**), these data collectively suggest that EBF1 impacts the distribution of cohesin on chromatin, with more pronounced effects at gene-sparse hubs.

**Figure 5:**
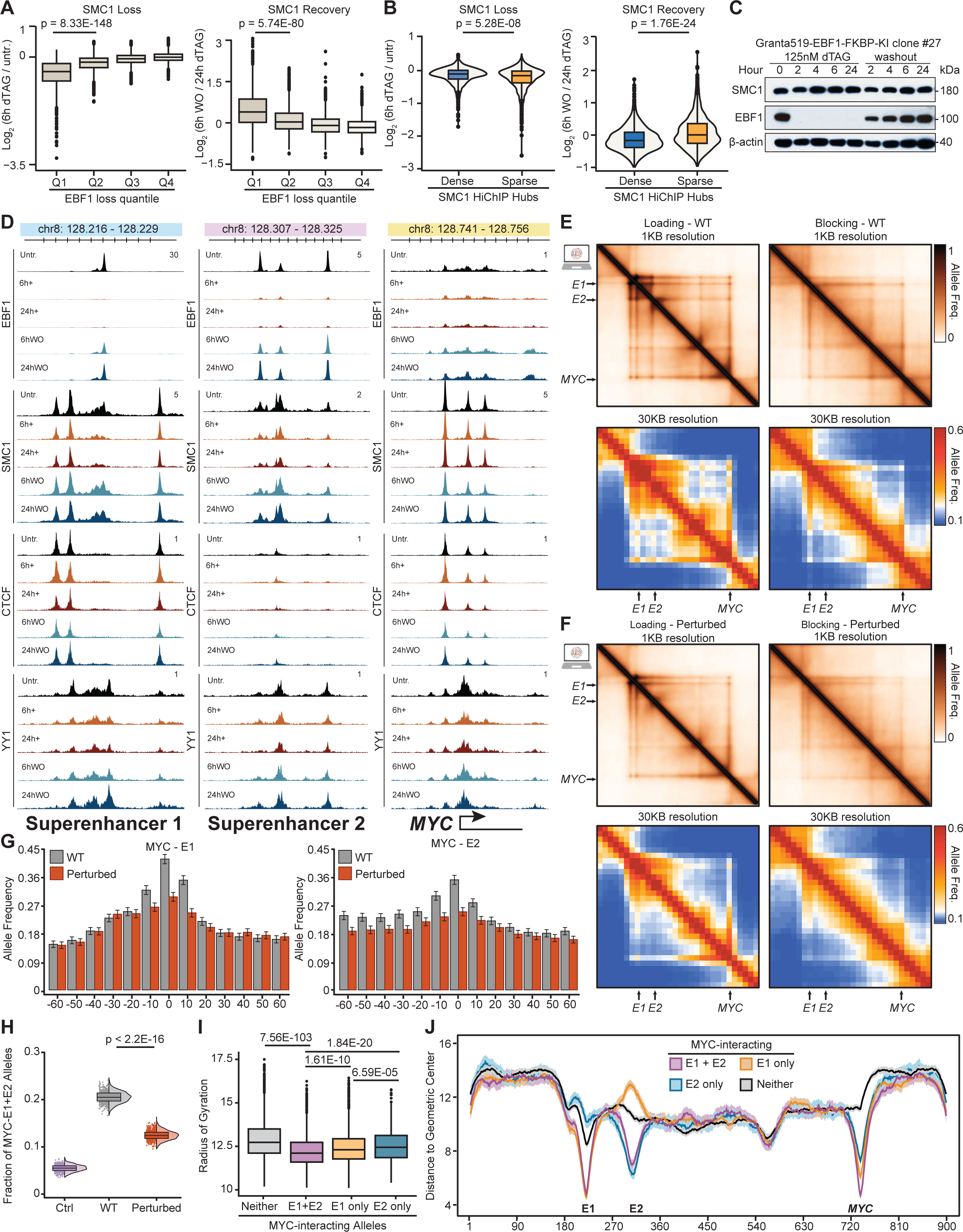
EBF1 positions enhancers to the *MYC* locus geometric centers by constraining cohesin traffic. A: Boxplots showing differential SMC1 ChIP-seq signals in 6h dTAG treatment (left) and washout (right) per quartile of EBF1 loading changes 6h after dTAG treatment. B: Box and violin plots comparing differential SMC1 ChIP-seq signals in 6h dTAG treatment (left) and washout (right) at SMC1 peaks located in gene-sparse and gene-dense hubs. P-values: Wilcoxon rank sum test. C: Time-course western blotting of SMC1 and EBF1 in Granta519 EBF1-FKBP-KI clone 27 showing SMC1 protein is invariant to EBF1 removal and restoration. ý-actin is loading control. D: ChIP-seq tracks showing EBF1, SMC1, CTCF1 and YY1 signals in MCL at the *MYC* super-enhancers E1 and E2 centered around EBF1 binding sites and the *MYC* gene body. Note concordant changes in SMC1 and EBF1 levels at the EBF1-bound sites after dTAG treatment and washout. E, F: Allele frequency maps for simulated *MYC* locus polymers in EBF1 wildtype (E) and EBF1- degraded (perturbed) (F) conditions. Allele frequencies are calculated using 10 as the cutoff for interaction between two monomers. The frequency of pairwise interactions between 900 monomers is equivalent to 1 Kb genomic resolution (top rows), and the median of pairwise interaction frequencies of each 30 monomers is equivalent to 30 Kb resolution (bottom rows). See methods for the parameters distinguishing EBF1 as an extruder loader (left column) or barrier (right column). G: Barplots of allele frequencies of monomers representing E1 (left), E2 (right) and *MYC* compared with the distance-matched pairs using a sliding window (see Methods). Error bars show standard deviation of allele frequency calculated from 1,000 randomly sampled polymers in 1,000 iterations. H: Box and violin plots showing the decrease of *MYC*-E1-E2 interaction frequencies in perturbed *MYC* locus. Each point represents the allele frequencies calculated from 1,000 randomly sample polymers with a total of 1,000 iterations per condition. Control regions are shifted 60 monomers to the left of *MYC*, E1 and E2 monomers. P-values: Wilcoxon rank sum test. I: Boxplots showing distribution of radius of gyration of simulated polymers with *MYC* interacting with E1, E2, both or neither in wildtype scenario. P-values: Wilcoxon rank sum test. J: Distances of each monomer to the geometric centers of the polymers with *MYC* interacting with E1, E2, both or neither. The median values and standard deviations across stratified polymers are calculated and plotted for each of the 900 monomers in wildtype scenario.

### EBF1 positions enhancers to the spatial center of *MYC* locus by constraining cohesin traffic in MCL

Considerable evidence supports the role of cohesin in shaping genome topology through the loop extrusion model^68-70^. After loading onto chromatin, cohesin complexes dynamically extrude chromatin until they are unloaded, or their movements are blocked and/or paused by extrusion barriers such as CTCF. To gain a more mechanistic understanding of the interplay between EBF1, cohesin and enhancer positioning in MCL, we conducted an in-depth analysis of SMC1 chromatin occupancy at 7,777 bona fide EBF1-bound elements (**Fig. S5C**), which revealed two distinct patterns of EBF1 binding: 1,536 sites with broad and 5,435 sites with focal EBF1 occupancy (**Fig. S5C**). High SMC1 levels were detectable at ∼90% of EBF1-bound elements (**Fig. S5C**), whereas CTCF was only detected at a small subset of focal EBF1-bound elements. Notably, we observed significantly greater decreases in SMC1 occupancy at the broad compared to focal EBF1-bound elements 6 hours after EBF1 removal (**Fig. S5D**), as exemplified by *NEK2, UST*, and *MYC* loci (**Figs. 5D, S5E and S5F**). Close investigation of broad EBF1 binding at these loci showed that EBF1 removal and restoration markedly changed SMC1, but not CTCF, levels both at the EBF1-bound elements and their immediate flanking sequences (**Figs. 5D, S5E and S5F**). Extending this analysis genome-wide showed that EBF1 loss decreased SMC1 occupancy at the flanking sequences of EBF1 binding sites, with a greater effect at broad EBF1-bound sites (**Fig. S5G**). Collectively, these data reinforce our earlier finding that SMC1 occupancy is EBF1-dependent (**Fig. 5A**) and further suggest a potential role for EBF1 in influencing cohesin movement on chromatin, leading to cohesin accumulation at and near EBF1-bound elements.

Led by our findings that EBF1 impacts both cohesin chromatin occupancy and *MYC* multi-enhancer-promoter interactions, we next sought to unify these observations and determine how EBF1-mediated cohesin redistribution contributes to enhancer positioning. To this end, we considered two hypotheses. First, EBF1 may increase the loading of cohesin onto chromatin, which in turn facilitates extrusion of EBF1-bound enhancers towards promoters. Alternatively, EBF1 might impede cohesin movement along the chromosome, which could also explain cohesin enrichment at and near EBF1-bound sites (**Figs. 5D, S5E and S5F**). In the second model, EBF1 slows and/or pauses cohesin’s typical translocation at its binding sites, thereby increasing the frequency of alleles with enhancer-promoter interactions.

To differentiate between these two hypotheses, we leveraged our ORCA data and physical polymer modeling to examine the molecular mechanism that best explains *MYC* locus topological structures and their EBF1 dependencies at both the population and single-chromatin fiber levels. In both *MYC* locus models, we considered 900-Kb-long synthetic polymers and introduced two strong extrusion barriers at the centromeric and telomeric TAD boundaries (**Fig. 5E and Table S7**). The sizes of barriers were set according to the genomic spans of CTCF- and cohesin-enriched elements at the *MYC* locus. We differentiated the two models based on EBF1’s relationship with parameters governing loop-extruding factor loading and movement.

In the first model, consistent with the role of EBF1 as a cohesin loading facilitator, extruding factors are loaded at specific polymer regions that correspond to EBF1-enriched elements in our ChIP-seq data. In contrast, in the second model, EBF1 binding define extrusion barriers rather than loading sites. Specifically, the EBF1-enriched elements determine the span and blocking capability of TAD-internal permeable barriers that obstructed the movement of randomly loaded extruding factors on polymers.

With the same density and lifetime of extruders, population-level maps of both models resulted in features reminiscent of architectural stripes, loops, and TAD boundaries (**Fig. 5E**). The model in which EBF1 acted as a cohesin loader exhibited intense enrichment of interactions near the loading sites, similar *MYC*-to-E1 and *MYC*-to-E2 interaction frequencies, and depletion of intra-TAD interactions, suggesting that this model favors the formation of short-range, point-to-point chromatin loops (**Fig. 5E**). In contrast, the model in which EBF1 acted as a loop-extrusion barrier more accurately captured the diverse population-level features observed at the *MYC* locus by sequencing and ORCA imaging assays (**Figs. 3D and 3E**), including the differing intensities of *MYC*-to-E1 and *MYC*-to-E2 looping, as well as intra-TAD interactions at evenly distributed frequencies (**Fig. 5E**).

Encouraged by these data, we further explored how these two plausible mechanisms could explain EBF1’s impact on pairwise *MYC* promoter-enhancer interactions. The loss of EBF1 as a extruders’ loader predicted a complete loss of intra-TAD structure, with the persistence of chromatin-jet-like interactions^71^ near cohesin loading sites – features that were not observed in sequencing or imaging of the *MYC* locus (**Figs. 5E, 5F, 3D and 3E**). By contrast, loss of EBF1 as a loop-extrusion barrier more precisely reproduced ORCA imaging data and demonstrated a more pronounced decrease in *MYC*-to-E2 compared with *MYC*-to-E1 loop and stripe features (**Figs. 5E, 5F and 3E**). This model also more closely recapitulated the impact of EBF1 on intra-TAD interactions observed at the *MYC* locus by ORCA imaging (**Figs. 5E, 5F and 3E**).

To further investigate the role of EBF1 as a barrier to cohesin movement, we complemented our physical polymer modeling studies by examining insulation potential of EBF1-bound sites using Micro-C data. We observed stronger insulation at the EBF1-bound sites compared to their flanking sequences. Notably, the loss of EBF1 markedly decreased insulation potential of these sites without affecting insulation at other accessible elements including CTCF-bound elements (**Figs. S5H and S5I**).

Given that the role of EBF1 as a loop-extrusion barrier more closely explained population-level features and EBF1-dependency of *MYC* locus topology (**Figs. 5E, 5F and 3E**), we next compared multi-enhancer-promoter frequency and enhancer central positioning observed in experimental and synthetic single chromatin fibers. Similar to experimental data (**Fig. 4E**), EBF1 loss decreased the frequency of polymers with *MYC*-E1-E2 three-way interactions (**Figs. 5G and 5H)**. Consistent with single-chromatin experimental data (**Fig. 4H**), the E1 and E2 regions were displaced from the geometrical centers in polymers lacking *MYC*-to-E1 and *MYC*-to-E2 interactions, respectively (**Figs. 5I and 5J**). Finally, these studies showed that in polymers with *MYC*-E1-E2 three-way interactions, all three segments were positioned closer to the polymer’s geometric centers (**Figs. 5I and 5J**), recapitulating ORCA data (**Fig. 4H**). Collectively, these analyses demonstrate that a constrained loop-extrusion mechanism more accurately explains our population- and single-cell-level experimental observations, suggesting that EBF1-mediated changes in cohesin traffic patterns at the *MYC* locus constrains enhancer positioning to organize *MYC* multi-enhancer interactions in MCL.

### EBF1 binds and activates *KLF7* promoter without changing its local radial positioning in GSI-resistant T-ALL

We next investigated the generalizability of our findings in MCL by extending our studies to T acute lymphoblastic leukemia (T-ALL), where EBF1 derepression is required for the acquisition of resistance to the Notch antagonist gamma-secretase inhibitor (GSI)^72^. Our MCL data showed that TADs and compartments remained invariant to acute short-term EBF1 changes (**Figs. 1E and S1J**). By contrast, our earlier studies in T-ALL revealed B-to-A compartment shifts in over 100 loci during long-term EBF1 derepression in GSI-resistance acquisition^72^.

The locus encompassing *KLF7*, a regulator of T cell proliferation^73,74^, was one of the loci that moved from B to A compartment (**Fig. S6A**). Concordantly, GSI-resistant cells exhibited high expression of *KLF7*, whereas the gene was inactive in GSI-sensitive cells^72^. Notably, H3K27ac and EBF1 chromatin occupancy analyses showed prominent gains in chromatin activity and EBF1 binding exclusively at the *KLF7* gene body, with no significant changes at any distal element (**Fig. S6A**). We thus leveraged the *KLF7* locus to investigate whether gene activation alone is sufficient for central positioning.

To examine this hypothesis, we first utilized Hi-C and Micro-C to map the population-average folding of the *KLF7* locus. These data showed reproducible increases in population-average interaction frequencies and TAD boundaries’ insulations across the *KLF7* locus in GSI-resistant T-ALL (**Figs. S6B and S6C**). We next performed 37-step ORCA experiments to investigate the impact of EBF1 on the organization of individual *KLF7* alleles in GSI-resistant T-ALL (**Fig. S6A**). Our 8,128 primary oligonucleotides (**Table S5**) spanned the B-to-A-compartment-switched region. In contrast to the *MYC* promoter in MCL, the *KLF7* promoter was approximately located at the midpoint of the traced locus on the linear genome (**Fig. S6A**). Imaging of ∼11,000 *KLF7* alleles in GSI-sensitive and GSI-resistant T-ALL complemented our population-average assays (**Figs. S6B and S6C**) and revealed marked decreases in pairwise distances between most segments of the *KLF7* locus during drug resistance acquisition (**Fig. S6D**). Furthermore, this single-allele data demonstrated significant compaction of the *KLF7* locus chromatin fiber in GSI-resistance (**Fig. S6E**).

We next leveraged the single-allele nature of our chromatin tracing data to examine the differential local radial positioning of genomic elements at the *KLF7* locus during GSI resistance acquisition. In GSI-sensitive cells, the 3’ TAD boundary (step 31) and several SMC1 and CTCF co-bound elements were the closest segments to the geometric centers of chromatin fibers (**Figs. S6F and S6G**). Similarly, analysis of GSI-resistant cells revealed central positioning of elements with a concomitant gain of CTCF and SMC1 (**Figs. S6F, S6G and S6A**), potentially facilitating the multiple stripe-like features observed in the GSI-resistant population-average interaction frequency map (**Fig. S6D**). More importantly, this analysis revealed that despite being near the locus’s midpoint on the linear genome (**Fig. S6A**), the local radial positioning of the *KLF7* promoter remained unchanged in GSI-resistance (**Fig. S6F**). Collectively, chromatin tracing of the *KLF7* locus during GSI-resistance acquisition demonstrated that gene activation is not sufficient for the central positioning of regulatory elements.

### EBF1 positions *ZEB2* and its enhancer to allelic spatial center to enable their interactions in GSI-resistant T-ALL

Similar to *KLF7*, the T-ALL-associated transcription factor *ZEB2*^75^ also shifted from B to A compartment and was derepressed during GSI-resistance development in T-ALL (**Fig. 6A**). However, unlike *KLF7*, EBF1 bound to and activated a distal enhancer ∼200 Kb 5’ of the *ZEB2* promoter (**Fig. 6A**). Notably, Hi-C and Micro-C analyses showed the formation of a stripe-like feature between the GSI-resistance-restricted EBF1-bound enhancer and *ZEB2,* but not *GTDC1*, which was already expressed in GSI-sensitive cells (**Figs. 6A-C**). Additionally, we observed increased SMC1 loading at the *ZEB2* promoter and its interacting EBF1-bound enhancer, but not at the *GTDC1* promoter (**Fig. 6A**). In sum, these data suggest that, in contrast to *KLF7*, the acquisition of an EBF1-bound enhancer contributes to *ZEB2* upregulation in GSI-resistance^72^, allowing us to investigate whether EBF1-dependent distal enhancer activation alters local radial positioning of enhancers and promoters.

**Figure 6:**
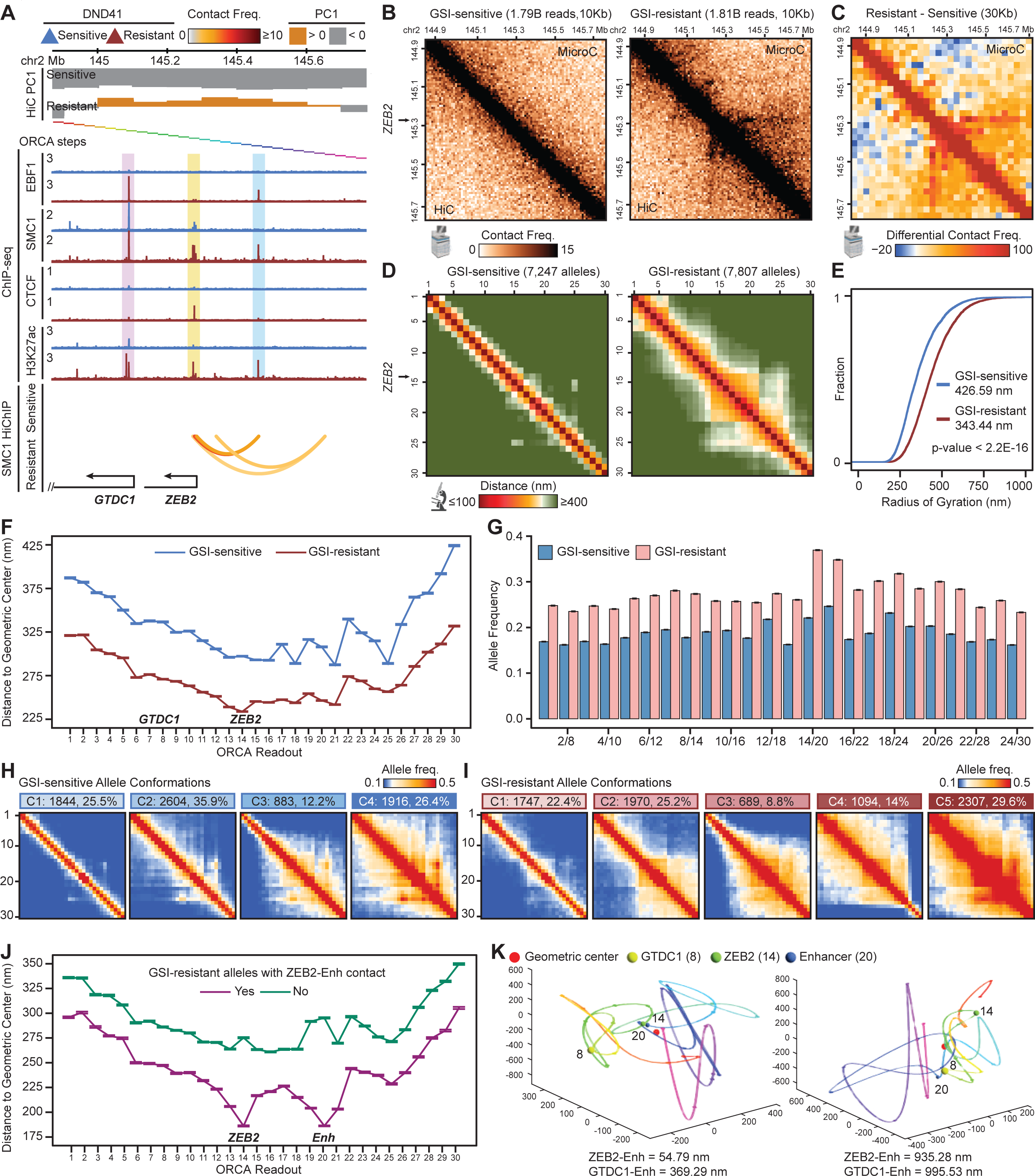
EBF1 positions the *ZEB2* promoter and enhancer towards the locus geometric centers in GSI-resistant T-ALL. A: ChIP-seq tracks showing gain of active histone marks H3K27ac, EBF1, CTCF and SMC1 at *ZEB2* locus in GSI-resistant DND41. Normalized SMC1 HiChIP arches showing gain of interactions in GSI-resistant DND41. The PC1 values of Hi-C contact correlation matrices in GSI- sensitive and GSI-resistant cells showing shift from B (<0) to A (>0) compartment. Rainbow-colored bars indicate the 30 steps of ORCA experiments. Bottom track indicates the positions of expressed genes. B: Normalized Micro-C (top) and Hi-C (bottom) interaction frequency maps showing gain of interactions in GSI-resistant (right) compared to GSI-sensitive (left) at *ZEB2* locus in GSI-sensitive vs GSI-resistant DND41. C: Differential (GSI-resistant minus GSI-sensitive) normalized Micro-C interaction maps with 30 Kb resolution matching ORCA showing gain of stripe between *ZEB2* promoter and enhancer in GSI-resistant DND41. D: ORCA pairwise distance maps of *ZEB2* chromatin traces in GSI-sensitive (left) and GSI- resistant (right) DND41. Each point represents the median of pairwise distances between two segments across all alleles. Alleles are not imputed, and missing values are excluded from the calculation. E: Cumulative distribution of radius of gyration of *ZEB2* chromatin traces in GSI-sensitive and GSI-resistant DND41 showing overall compaction of *ZEB2* locus in GSI-resistant cells. P-value: Kolmogorov–Smirnov test. F: Distances of each segment to the geometric centers of the chromatin traces of *ZEB2* locus in GSI-sensitive and GSI-resistant DND41. The median values of chromatin traces are calculated and plotted for each ORCA segment in GSI-sensitive and GSI-resistant cells. Error bars show 95% confidence interval from bootstrapping. Note that *ZEB2* promoter (step 14) but not *GTDC1* (step 8) is positioned the closest to geometric centers of alleles in GSI-resistance. G: Barplots of allele frequencies with *ZEB2* promoter-enhancer interactions (steps 14 and 20) in GSI-sensitive and GSI-resistant DND41 compared with the distance-matched pairs using a sliding window (see Methods). Cutoff for interaction: 250nm. Bootstrapping is performed per condition and error bars show 95% confidence intervals. H, I: Allele frequency maps of chromatin traces for each of the 5 common topological conformations of the *ZEB2* locus in GSI-sensitive (H) and GSI-resistant (I) DND41. Clustering and calculation of frequencies are separately performed on pairwise distance matrices of alleles from each condition. J: Distances of each segment to the geometric centers of chromatin traces with or without *ZEB2* promoter-enhancer interactions in GSI-resistant DND41. The median values of stratified alleles are calculated and plotted for each ORCA segment. Error bars show 95% confidence interval from bootstrapping. K: Example reconstructed chromatin traces for alleles in GSI-resistant DND41 with (left) and without (right) *ZEB2* enhancer-promoter interaction emphasizing the central positioning of interacting elements.

To examine the impact of EBF1 on the local chromatin folding of individual *ZEB2* alleles, we performed 30-step ORCA experiments in GSI-sensitive and GSI-resistant T-ALL (**Fig. 6A**). Following the *KLF7* setup, we designed 5,912 primary oligonucleotide probes to trace the region that switched from B to A compartment (**Table S5**), with the *ZEB2* promoter located at the traced chromatin midpoint on the linear genome (step 14).

Similar to *KLF7*, we observed marked increases in average interaction frequencies (**Fig. 6B**), decreases in pairwise spatial distances (**Fig. 6D**), and significant locus compaction (**Fig. 6E**) in GSI resistance. Despite these similarities, the *KLF7* and *ZEB2* loci exhibited distinct changes in local radial positioning. While the centrality of the *KLF7* promoter remained unchanged, the *ZEB2* promoter became the closest segment to the locus’s geometric center in GSI resistance (**Fig. 6F**). Notably, the *ZEB2* promoter and its GSI-resistance-restricted EBF1-bound enhancer (step 20) exhibited the highest increase in interaction frequencies compared to distance-matched internal controls (**Fig. 6G**). Examination of high-dimensional chromatin traces further revealed that, unlike the short-term EBF1 impact on MCL topology, prolonged EBF1 expression established new common topological conformations at the *ZEB2* locus during GSI-resistance development (**Figs. 6H and 6I**). Interestingly, many of these newly formed conformations were permissive to *ZEB2* promoter-enhancer interaction.

Encouraged by these observations, we compared centrality profiles of traces with and without *ZEB2* promoter-enhancer interaction in GSI-resistance to assess the role of local radial positioning in promoter-enhancer interactions. Importantly, we found that the *ZEB2* promoter and its EBF1-bound enhancer were markedly placed towards the geometric center relative to the rest of the locus only in traces where the enhancer-promoter interaction was present (**Figs. 6J and 6K**). By contrast, traces from GSI-resistant T-ALL lacking *ZEB2* promoter-enhancer interactions exhibited a centrality pattern similar to that of GSI-sensitive cells (**Figs. 6J and 6F**). Together, our single-allele-resolution study of *ZEB2* and *KLF7* chromatin topology in GSI-resistant T-ALL supports our observations in MCL and demonstrates that gene activation alone is not sufficient, but rather EBF1-dependent distal enhancer activation is required for central positioning.

### EBF1-mediated cohesin redistribution in MCL is also observed in GSI-resistant T-ALL

Intrigued by the finding that EBF1-instructed radial positioning of enhancers and promoters extends to GSI-resistant T-ALL, we next examined whether the EBF1-mediated changes in cohesin traffic pattern observed in MCL were also generalizable to T-ALL. We identified 17,602 EBF1-bound elements in GSI-resistant cells, 78.5% of which exhibited marked SMC1 occupancy (**Fig. S6H**). Close examination of these elements revealed several distinct patterns of EBF1 and SMC1 co-occupancy (**Fig. S6H**), exemplified by *ZNF195* and *GADD45B* loci (**Fig. S6I**). SMC1 occupancy paralleled chromatin accessibility at EBF1-bound elements, again suggesting a potential link between EBF1 chromatin binding and cohesin extrusion (**Figs. S6H and S6I**). Similar to MCL (**Fig. S5C**), CTCF was detected at a small subset of EBF1-bound elements in GSI-resistant T-ALL (**Fig. S6H**). Despite exhibiting the strongest insulation potential (**Fig. S6K**), these elements showed the lowest SMC1 gain in GSI resistance (**Fig. S6J**). A comparison of GSI-sensitive and -resistant T-ALL further demonstrated that, similar to the broad EBF1-bound elements in MCL (**Fig. S5C**), the most significant SMC1 gains occurred at the strongest EBF1- bound elements lacking CTCF binding in GSI-resistance (**Figs. S6H and S6J**). Finally, we observed stronger insulation potential at EBF1-bound elements compared to their flanking sequences in T-ALL (**Fig. S6K**). Together, these data suggest that similar to MCL, gain of EBF1 binding during GSI-resistance acquisition alters cohesin traffic pattern in T-ALL.

### The *MYC* promoter and enhancers interact at the allelic spatial centers in individual GSI-sensitive and GSI-resistant T-ALL cells

Our data so far suggest that EBF1, a crucial transcription factor for MCL and GSI-resistant T-ALL, facilitates the positioning of distal enhancers and promoters to the geometric centers of loci to promote their interactions. We next asked whether other transcription factors similarly impact the local radial positioning of these regulatory elements. To test this, we hypothesized that T-cell-specific transcription factors, including TCF1^12^, position active enhancers at allelic geometric centers to facilitate their interactions with target gene promoters in GSI-sensitive T-ALL, where EBF1 is not expressed. More specifically, we focused on T-ALL-restricted super-enhancers TE1 and TE2, located 1.35 Mb and 1.85 Mb 3’ of the *MYC* promoter in GSI-sensitive T-ALL, respectively (**Figs. 7A and 3A**). We previously demonstrated that differential activity of TE1 and TE2 maintains *MYC* expression during GSI-resistance acquisition in the absence of any genetic mutation^72^.

**Figure 7:**
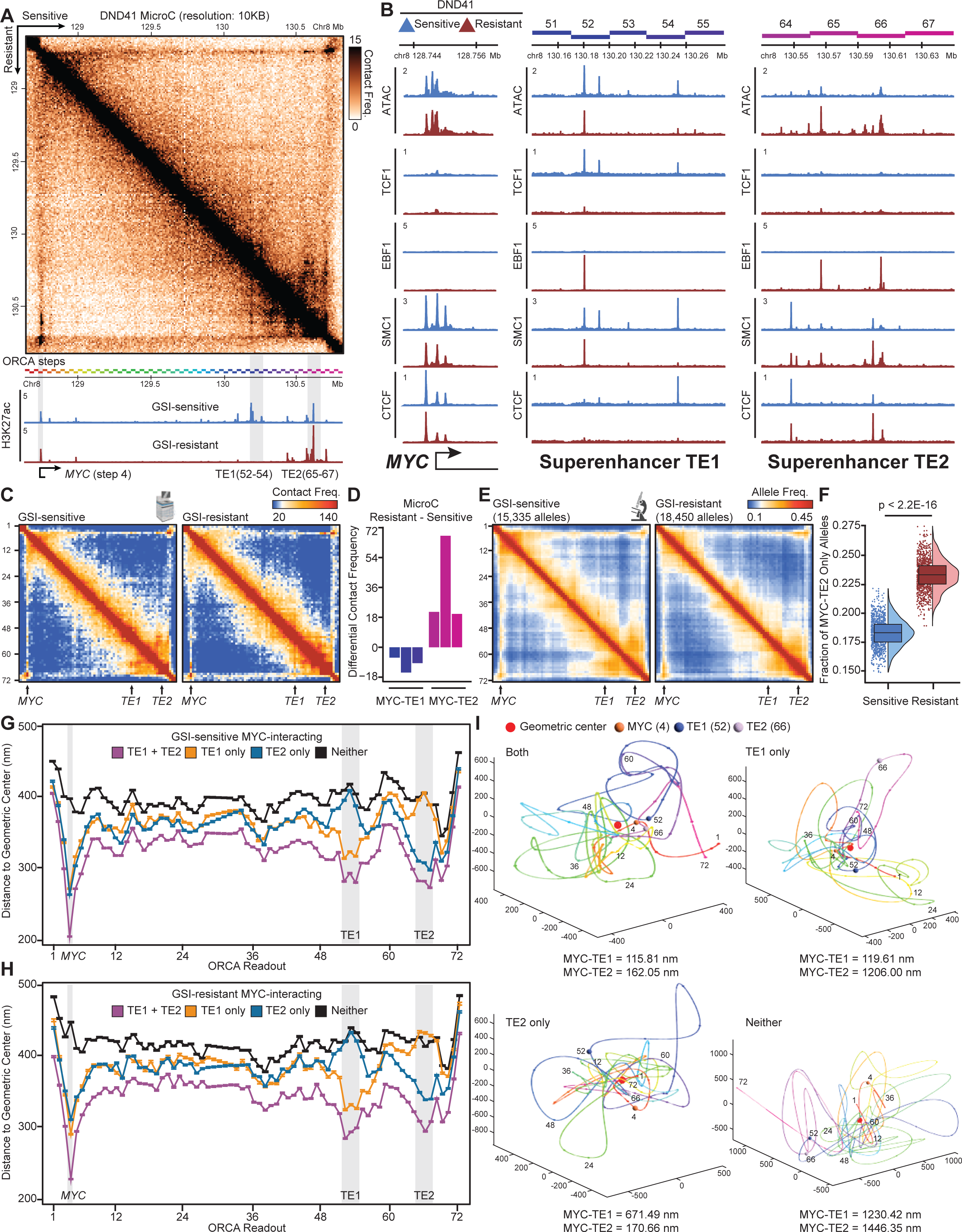
Two *MYC* distal super-enhancers interact with the promoter near allelic geometric centers in GSI-sensitive and GSI-resistant DND41. A: Top: Normalized Micro-C contact maps at 10 Kb resolution in GSI-sensitive (top) and GSI- resistant (bottom) DND41. Bottom: H3K27ac ChIP-seq tracks showing two super-enhancers TE1 and TE2 located more than 1.5 Mb 3’ of the *MYC* promoter in GSI-sensitive DND41. Note loss of TE1 and increase in TE2 activity in GSI-resistant DND41. Rainbow-colored bars indicate the 72 steps of ORCA experiments. B: ATAC-seq and ChIP-seq genome tracks showing invariant accessibility, SMC1 and CTCF levels at the *MYC* promoter in GSI-sensitive and GSI-resistant DND41. In contrast, TCF1-bound super-enhancer TE1 lost most of its accessible elements, as well as SMC1 and CTCF binding while super-enhancer TE2 gained EBF1, SMC1 and CTCF binding and accessibility at several elements. Colored bars on the top indicate coverage of ORCA steps. C: Normalized Micro-C interaction frequency maps in GSI-sensitive (left) and GSI-resistant (right) DND41 at 30 Kb resolution matching ORCA steps showing a stripe-like feature connecting the *MYC* promoter and the two distal super-enhancers. D: Quantification of Micro-C normalized interaction frequencies between the *MYC* promoter (bin number 4) and TE1 (bin numbers 52-54) or TE2 (bin numbers 65-67) in GSI-sensitive and GSI- resistant DND41. E: Allele frequency maps for GSI-sensitive (left) and GSI-resistant (right) DND41 *MYC* chromatin traces from two biological replicates per condition. Each condition uses the average distance between two consecutive segments as the cutoff for determining interaction. Alleles are not imputed, and missing values are excluded from the calculation of frequencies. F: Box and violin plots showing increase in allele frequency with only *MYC*-TE2 interaction in GSI- resistant DND41. Each point represents the allele frequencies calculated from 1000 randomly sampled alleles with a total of 1000 rounds of random sampling per condition. P-values: Wilcoxon rank sum test. G, H: Distances of each segment to the geometric centers of chromatin traces with *MYC*-TE1- TE2, only *MYC*-TE1, only *MYC*-TE2 or no *MYC* promoter-enhancer interactions (Neither) in GSI- sensitive (G) and GSI-resistant (H) DND41. The median values of stratified alleles are calculated and plotted for each ORCA segment in untreated cells. Error bars show 95% confidence interval from bootstrapping. I: Example reconstructed chromatin traces for alleles with *MYC*-TE1-TE2 three-way interaction or only *MYC*-TE1 interaction from GSI-sensitive DND41, and only *MYC*-TE2 interaction or no promoter-enhancer interactions (Neither) from GSI-resistant DND41 emphasizing the central positioning of interacting elements.

Close examination of the super-enhancer TE1 in GSI-sensitive cells identified three TCF1-bound accessible elements with high levels of H3K27ac and SMC1 (**Figs. 7A and 7B**). While all three elements were deacetylated, only two of them became inaccessible, while the third remained accessible in GSI-resistant cells. Notably, we observed the loss of TCF1 and the gain of EBF1 binding at the single TE1 element that remained accessible and SMC1-occupied in both GSI-sensitive and resistant cells. Our high resolution and unbiased Micro-C analysis corroborated our previous Hi-C, SMC1 HiChIP and UMI-4Cseq data^72^, showing that the *MYC* promoter interacted with the TE1 super-enhancer - albeit at a lower frequency - in GSI-resistant T-ALL, potentially due to replacement of TCF1 with EBF1 (**Figs. 7A, 7C and 7D**).

Similar to super-enhancer TE1, our detailed study of super-enhancer TE2 in GSI-sensitive T-ALL identified several highly acetylated and accessible elements (**Figs. 7A and 7B**). These elements did not bind EBF1 or TCF1 and were potentially regulated by other transcription factors in GSI-sensitive T-ALL (**Figs. 7A and 7B**). In contrast to TE1, several elements in TE2 super-enhancer bound EBF1 and concomitantly gained marked accessibility, acetylation, and SMC1 in GSI- resistance (**Figs. 7A and 7B**). Consistent with these changes, we observed an increase in the population-average *MYC*-to-TE2 interaction frequency in GSI-resistance (**Figs. 7A, 7C and 7D**). Together, these data suggest that differential binding of lineage-specific transcription factors alters the activity, cohesin loading, and positioning of the two T-ALL-restricted *MYC* super-enhancers during GSI-resistance acquisition.

We next performed 72-setp ORCA experiments to trace the folding of individual *MYC* chromatin fibers in GSI-sensitive and GSI-resistant T-ALL (**Table S5**), where different lineage-determining transcription factors bound super-enhancers TE1 and TE2 (**Fig. 7A**). Given the marked reproducibility, stability (**Figs. S7A and S7B**), and hybridization efficiency (**Fig. S7C** and **Table S6**), we combined biological duplicates and traced a 2.16-Mb chromatin region 3’ of *MYC* in 15,335 GSI-sensitive and 18,450 GSI-resistant alleles. In both conditions, the population-average ORCA and Micro-C-detected pairwise interaction maps showed a strong correlation (**Figs. 7C, 7E, and S7D**). Furthermore, the *MYC* TAD boundaries, enhancer-promoter loop dots, and stripe-like feature connecting the *MYC* promoter, TE1 and TE2 were clearly observed in ORCA’s average distance and interaction frequency maps, matching those identified by Micro-C (**Figs. 7C, 7E, and S7D**).

Ensured by the high quality of our ORCA data, we sought to investigate the impact of transcription factor binding on the positioning of super-enhancers TE1 and TE2 in GSI-sensitive and GSI-resistant T-ALL. To this end, we first examined pairwise distances between ORCA probe sets overlapping with the *MYC* promoter (step 4), super-enhancer TE1 (steps 52-54), and super-enhancer TE2 (step 65-67). The *MYC* promoter was separated from the TE1 in GSI-resistant cells (**Fig. S7E**). Conversely, TE2 were closer to the *MYC* promoter in resistant T-ALL (**Fig. S7E**). Consistent with these observations, we found a decrease and an increase in the frequency of alleles with only *MYC*-to-TE1 and *MYC*-to-TE2 interactions in GSI-resistance, respectively (**Figs. 7F, S7F and S7G)**. In concordance with our observations in MCL, three-way interactions among *MYC* and the two distal super-enhancers were also present in T-ALL. Furthermore, we observed a mild but statistically significant decrease in the allele frequency of *MYC*-TE1-TE2 interactions in GSI-resistance (**Figs. S7F and S7G)**. Combined with our chromatin binding data (**Figs. 7A and 7B**), this analysis suggests that T-cell-specific transcription factors such as TCF1 position *MYC* distal enhancers in GSI-sensitive cells. By contrast, EBF1 derepression and chromatin binding positioned both TE1 and TE2 to the *MYC* promoter, even in the absence of TE1 enhancer activity.

Our optical tracing of individual *MYC* alleles in GSI-sensitive and resistant T-ALL so far confirmed our findings in MCL and further showed that EBF1 and TCF1 binding position both *MYC* enhancers regardless of their activity. We thus asked whether *MYC* and its two super-enhancers also interact at geometric centers of individual *MYC* alleles in GSI-sensitive and -resistant T-ALL, where the activity and genomic location of enhancers are completely distinct from those in MCL (**Fig. 3A**). T-ALL traces with *MYC*-TE1-TE2 three-way interactions were markedly more compact than those with only *MYC*-to-TE1, only *MYC*-to-TE2, or no *MYC* promoter-enhancer interactions (**Fig. S7H**). More importantly, examination of the traces containing *MYC*-TE1-TE2 interactions in GSI-sensitive cells revealed that *MYC*, TE1 and TE2, despite being at the two ends of a ∼2 Mb locus on the linear genome, were the closest segments to the alleles’ geometric centers (**Figs. 7G, 7I and S7I**). In contrast, segments of traces with no enhancer-promoter interactions in GSI-sensitive T-ALL had uniformly larger distances to the allelic geometric centers (**Figs. 7G, 7I and S7I**). We also observed marked separation of TE1 and TE2 from the loci centers in traces with only *MYC*-to-TE1 or *MYC*-to-TE2 interactions in GSI-sensitive T-ALL, respectively (**Figs. 7G, 7I and S7I**).

Extending this analysis to GSI-resistant T-ALL confirmed our observation in MCL and GSI-sensitive T-ALL and further showed that enhancers and promoters were positioned at the allelic geometric centers to interact (**Figs. 7H, 7I and S7I**). Notably, despite the lack of acetylation, we observed a high level of SMC1 at the EBF1-bound element of TE1 in GSI-resistant T-ALL (**Figs. 7A and 7B**). Interestingly, this element was the most proximal segment to the geometric centers of traces from GSI-resistant cells (**Fig. 7H**). Overall, these results confirm our earlier data in MCL and suggest that the central positioning of enhancers is a general mechanism leveraged by lineage-determining transcription factors such as EBF1 and TCF1 to facilitate multi-enhancer regulatory interactions.

## DISCUSSION

The prevalence and mechanisms by which multiple enhancers regulate a single target gene have long intrigued biologists^9,10,76-79^. In this study, we utilized rapid degradation of EBF1 in MCL as a model to investigate direct role of lineage-determining transcription factors in chromatin folding and multiway enhancer-promoter interactions. We found that EBF1 preferentially promotes the formation of chromatin loops connecting multiple distal enhancers and key B-cell genes, which are sparsely distributed across large genomic distances. Our time-resolved chromatin tracing showed heterogenous positioning of EBF1-bound enhancers in hubs and their dynamic responses to EBF1 modulation. We found that when alleles form enhancer-promoter interactions, EBF1 positions these elements at their allelic geometric centers. We demonstrated that these observations are generalizable to T-ALL and may also extend to other lineage-determining transcription factors, such as TCF1. We provided clear evidence that multiple enhancers and promoters can interact at individual alleles, preferentially located close to the geometric centers of local chromatin. Together these findings reveal a general mechanism by which lineage-determining transcription factors facilitate interactions of regulatory elements in nuclear space.

Enhancer-promoter interactions are being measured at an unprecedented scale and depth by chromatin conformation capture techniques. Micro-C, which uses micrococcal nuclease to digest chromatin and HiChIP, which uses ChIP reactions to enrich for chromatin-bound proteins, have significantly advanced our genome-wide understanding of enhancer-promoter interaction networks^22,23,26,27,40,42^. At selected genomic loci, primer-based methods with multiple replicates such as UMI-4C-seq allow robust statistical comparison of enhancer-promoter contact frequency changes^72,80^, addressing a common limitation in genome-wide assays. Other enrichment-based methods such as Region Capture Micro-C allow discovery of interactions between ultra-close enhancers and promoters^81^. However, despite the high resolutions of these methods, they fundamentally rely on proximity-based ligation of two DNA fragments and thus are limited to providing only pairwise measurements of interaction frequencies. Super-resolution imaging techniques, such as stochastic optical reconstruction microscopy (STORM) using renewable fluorophore-conjugated oligos, enable the visualization of chromatin structures with unprecedented detail^82-84^. However, these techniques often struggle to distinguish precise chromatin segments and linear genome sequences, rendering them less suitable for studying the spatial relationship among enhancers and promoters. Our study leveraged the advantages of ORCA to analyze over 100,000 alleles in regions exemplifying compartment switches, architectural stripe formation and multiway enhancer-promoter interactions in response to modulation of lineage-specific transcription factors. We demonstrated the extensive potential of ORCA in elucidating complex chromatin architectures, and thus expect application of this approach to a broad range of loci will elucidate additional mechanisms of enhancer-promoter interaction. Furthermore, we anticipate that combining ORCA with other imaging modalities will enable further exploration of structure-function relationships between multiway enhancer-promoter interactions, transcription, and phase-separated^85^ protein condensates in individual cells.

Our studies using rapid degradation of EBF1 in MCL and long-term EBF1 activation in drug-resistant T-ALL further demonstrated the cooperation between EBF1 and SMC1. Our data suggest that the EBF1-cohesin axis may also play similar roles in other cancer types related to EBF1 mutations or dependencies. Future investigations across various cancer and developmental systems will reveal whether additional lineage-determining transcription factors can form chromatin loops directly, load architectural proteins, obstruct loop extrusion, or participate in the regulation of multiway interactions via other novel mechanisms. Many lineage-determining transcription factors including EBF1 contain intrinsically disordered domains (IDR)^86-88^. These unstructured transcription factor domains are critical for recruitment of coactivators and the formation of liquid-like condensates. Specifically, the C-terminal IDR of EBF1 is required for the recruitment of the chromatin remodeler BRG1 to promote chromatin accessibility during normal B cell development^38^. Additionally, a subdomain of the EBF1 IDR interacts with members of the FET RNA-binding protein family to mediate phase separation. However, it remains unclear whether the EBF1 IDR is also responsible for its cooperation with cohesin, or whether EBF1’s abilities to promote multiway interactions and form condensates are related. More broadly, while earlier studies point to potential links between phase-separated droplets and multi-enhancer clusters, future studies are necessary to rigorously substantiate this hypothesis and unravel the interplay between these two phenomena. We anticipate that further examination of IDR-containing lineage-determining transcription factors will reveal both shared or distinct roles of these factors in mediating chromatin looping and condensate formation in health and disease.

## Authors Contributions

Conceptualization: Y.Z., R.B.F.; Methodology: Y.Z., A.J., G.V., R.B.F.; Investigation: Y.Z., N.B., R.G., G.V., R.B.F.; Formal Analysis: Y.Z., N.B., T.F., S.Y., R.B.F.; Resources and Reagents: R.G., J.A., G.N., G.V., R.B.F.; Writing-Original Draft: Y.Z., R.B.F.; Writing-Review & Editing: Y.Z., T.F., N.B., R.G., G.V., R.B.F.; Funding Acquisition: R.B.F., G.V.; Supervision: R.B.F., G.V.

## Acknowledgment

The authors thank Warren Pear, Junwei Shi, Nikolay Zolotarev and Charles Antony for their helpful discussions and technical advice. We are grateful to Katherine Lupo for providing critical technical support. We also thank Shelley Berger for generous support of Penn Epigenetics Institute. This work was supported by NIH grants U01-DK112217, R01-HL145754, R0-AI168240, U01-DK127768 (to G.V.); and R01-CA-248041, R01-CA-230800 (to R.B.F.).

## METHODS

### Lead contact

Further information and request for reagents may be directed to and will be fulfilled by the lead contact, Robert B. Faryabi.

### Materials availability

Plasmids and cell lines generated in this study will be available upon request.

### Data and code availability

- Next-generation sequencing data have been deposited at GEO and will be publicly available as of the date of publication. Accession number is GSE293368. Microscopy data reported in this paper will be deposited at the 4DN and OLIVE data portals and also shared by the lead contact upon request.
- This paper does not report any original code.
- Any additional information required to reanalyze the data reported in this paper is available from the lead contact upon request.

## EXPERIMENTAL MODEL AND SUBJECT DETAILS

### Cell culture

DND41, JVM-2 and PGA-1 were purchased from Leibniz-Institute DSMZ-German Collection of Microorganisms and Cell Lines (DSMZ, cat# ACC525, ACC12, ACC766). HEK293T (CRL-11268) was purchased from ATCC. Granta519 was from the Broad Novartis Cancer Cell Line encyclopedia^63^. DND41, JVM-2 and PGA-1 were cultured in RPMI 1640 (Corning, cat# 10-040- CM) supplemented with 10% fetal bovine serum (Hyclone, cat# SH30070.03), 2 mM L-glutamine (Corning, cat# 25-005-CI), 100 U/mL and 100 μg/mL penicillin/streptomycin (Corning, cat# 30- 002-CI), 100 mM nonessential amino acids (Gibco, cat# 11140-050), 1 mM sodium pyruvate (Gibco, cat#11360-070) and 0.1 mM of 2-mercaptoethanol (Sigma, cat# M3148). HEK293T and Granta519 were cultured in DMEM (Corning, cat# 10-013-CV) supplemented with 10% fetal bovine serum (Hyclone, cat# SH30070.03) and 100 U/mL and 100 μg/mL penicillin/streptomycin (Corning, cat# 30-002-CI). GSI-resistant DND41 cells were generated as previously described^72^. All cell lines, including the cell lines described below, were grown at 37 °C and 5% CO_2_ and were used at a low passage number (<12) and subjected to regular mycoplasma tests and short tandem repeat (STR) profiling.

### Lentiviral packaging

Lentivirus was produced in HEK293T cells as previously described^27^. Briefly, 4.5x10^6^ HEK293T cells were plated in 8 mL DMEM media in 10 cm dishes 12-16 hours before transfection. The lentiviral constructs, packaging plasmid (pCMVdelta) and envelope plasmid (VSV-G) were co-transfected using FuGene HD (Promega, cat# E2311). The cells were returned to the incubator for 6-8 hours before replacement with 6 mL media. Lentiviral supernatants were harvested 48 hours post-transfection, subjected to 0.45 μm filtration and stored at -80 °C.

### CRISPR-Cas9 Editing

CRISPR/Cas9 system was used for knocking out EBF1 in Granta519, JVM-2 and PGA-1. Codon-optimized version of Cas9 carrying puromycin resistance gene (Cas9-puro) and sgRNA vectors carrying mCherry (LRmCherry2.1)^89^ were used. Cells were transduced with Cas9-puro lentivirus by spinfection at 2000 rpm for 90 min at 22°C in the presence of 6 μg/mL polybrene (Sigma-Aldrich, cat# H9268). Transduced cells were selected 3 days after spinfection with incremental 0.5-2 μg/mL puromycin until most cells are viable and maintained in 1 μg/mL puromycin. Expression of Cas9 was confirmed with western blot (CST cat# 14697S). sgRNA targeting *EBF1* exon 3 (EBF1-g7) was optimized as previously described^72^. Cells transduced with sgRNA were sorted with DAPI- mCherry+ gating on FACS Aria and used for RNA-seq, Hi-C and apoptosis experiments in **Fig. 1**.

### Generation of Granta519 single cell clones with endogenous EBF1 tagged with FKBP12^F36V^ (EBF1-FKBP-KI)

sgRNA targeting the stop codon of human *EBF1* gene (EBF1-KI-gRNA) was designed with Benchling (https://www.benchling.com/) and cloned in lentiviral sgRNA vector expressing GFP (LRG). Repair template with FKBP12^F36V^-HA-HA-P2A-BSD cassette flanked by 447bp sequences homologous to upstream and downstream of EBF1 stop codon were designed with SnapGene and synthesized in pUC57 plasmid by GenScript (pUC57-FKBP). Granta519-Cas9-expressing cells were electroporated with LRG-EBF1-KI-gRNA and pUC57-FKBP using Invitrogen Neon™ Transfection System per manufacturer’s recommendations. Cells were replated in 12-well plate with antibiotics-free media for 24 hours and transferred to full media with blasticitin selection for 15 days. Live single cells were sorted to 96-well plates containing blasticitin and cultured till visible clones were formed and further expanded. Genomic DNA was extracted with Quick-DNA Miniprep Plus Kit 200 Preps (Zymo cat# D4069) and PCR primers targeting the insertion site were used to determine homozygous insertion of FKBP cassette. Single cell clones with homozygous insertion of FKBP cassette were subject to dTAG^V^-1 (Bio-Techne cat# 6914) dosage and time titration by western blot. Finally, several single cell clones including Granta519-EBF1-FKBP-KI 26, 27, 29 and 97 showed complete degradation of EBF1 in 24-hour 125 nM dTAG^V^-1 treatment and were further validated with time-course RNA-seq.

### RNA sequencing (RNA-Seq)

Strand-specific RNA-seq for DND41 GSI-sensitive and GSI-resistant cells were previously published (GSE173872)^72^. RNA-seq was performed in the following cells with three replicates per condition: Granta519-Cas9 control and LRmCherry2.1-EBF1-g7 cells sorted 3 days post transduction; Granta519-EBF1-FKBP-KI clones 26, 27, 29 and 97 treated with 125nM dTAG^V^-1 for 0, 6 and 24 hours; Granta519-EBF1-FKBP-KI clones 27 and 97 treated with 125nM dTAG^V^-1 for 24 hours and washed with dTAG-free media 5 times and replated for 6 and 24 hours (washout). In all experiments, 3-5x10^5^ cells were washed with 1X PBS and lysed with 350 μL RLT Plus buffer (Qiagen) supplemented with 2-mercaptoethanol, vortexed briefly and immediately proceeded to RNA isolation with RNeasy Plus Micro Kit (Qiagen cat# 74034). RNA integrity numbers were determined using TapeStation 4150 (Agilent), and all samples used for RNA-seq library preparation had RIN numbers greater than 9.5. 300-800 ng of total RNA was used, and libraries were prepared using the SMARTer Stranded Total RNA Sample Prep Kit-HI Mammalian (Clontech cat# 634873). Libraries were paired-end sequenced on Illumina NextSeq 550 (38bp+38bp) or Novaseq 6000 (61bp+61bp).

### Assay for Transposase-Accessible Chromatin Sequencing (ATAC-seq)

ATAC-seq was performed as previously described^90^ and three replicates were performed for each condition. Briefly, 5x10^4^ cells were pelleted at 800 G and washed with 50 μL of ice cold 1X PBS (Corning cat# 21031CV), followed by 2 min treatment with 50 μL lysis buffer (10 mM Tris-HCl, pH 7.4, 3 mM MgCl_2_, 10 mM NaCl, 0.1% Igepal cat# CA-630). Pelleted nuclei were resuspended in 50 μL of transposition buffer (25 μl of 2X TD buffer, 22.5 μL of molecular biology grade water and 2.5 μL Tn5 transposase (Illumina cat# FC-121-1030)) to tag the accessible chromatin for 45 min at 37 °C. Tagmented DNA was purified with MinElute Reaction Cleanup kit (Qiagen cat# 28204) and amplified with 7 PCR cycles. Libraries were purified using QiaQuick PCR purification kit (Qiagen cat# 28106) and eluted in 20 μL EB buffer. Indexed libraries were assessed for nucleosome patterning on TapeStation 4150 (Agilent) and paired-end sequenced on Illumina NextSeq 550 (38bp+38bp) or Novaseq 6000 (61bp+61bp).

### Chromatin Immunoprecipitation Sequencing (ChIP-seq)

ChIP-seq for histone marks was performed as previously described^27^. Briefly, 1x10^7^ cells were crosslinked with 1% formaldehyde and lysed. Nuclei were extracted and resuspended in 1% SDS and sonicated using Brandson 450 sonicator with 25% amplitude, 0.5 sec on, 1 sec off for a total of 4 min 30 sec on. Solubilized chromatin was then diluted and cleared with IgG (CST cat# 2729S) and recombinant protein G–conjugated Agarose beads (Invitrogen cat# 15920-010) for 1 hour at 4°C. The cleared supernatant was subsequently immunoprecipitated with antibodies recognizing H3K27ac (Active Motif cat# 39133), H3K4me1 (Abcam cat# ab8895) or H3K27me3 (CST cat# 9733) overnight at 4°C. Buffers in all steps above were supplemented with protease inhibitors (Roche cat# 11697498001). Antibody-chromatin complexes were captured with recombinant protein G–conjugated Agarose beads, washed with low salt wash buffer, high salt wash buffer, LiCl wash buffer and TE buffer with 50 mM NaCl and eluted. Input sample was prepared by the same approach without immunoprecipitation.

ChIP-seq for transcription factors was performed as previously described^91^. Briefly, Dynabeads Protein G (ThermoFisherScientific cat# 10003D) was incubated with the following antibodies: 1) polyclonal anti-EBF1 (1C) antibody, which recognizes an N-terminal EBF1 peptide (RG)^38^, 2) commercial EBF1 antibody (Millipore cat# AB10523), 3) SMC1a (Bethyl, cat# A300-055A), YY1 (Active motif cat# 61779) and CTCF (EMD Millipore cat# 07-729) for 8-12 hours in PBS+0.5% BSA at 4 °C. 4x10^7^ cells were crosslinked with 1% formaldehyde and 1.5 mM EGS (ThermoFisherScientific cat# 21565) and sonicated using Brandson 450 sonicator with 17% amplitude, 10 sec on, 1 min off for 10 times. Lysate was then cleared by centrifuging for 5 min at 16 Kg, 4 °C and incubated with antibody-bound beads and 1% Triton Tx-100 (Roche cat# 10789704001) overnight at 4 °C. Buffers in all steps above were supplemented with protease inhibitors (Roche cat# 11697498001). Antibody-chromatin complexes captured on beads were then separated on magnet and washed with wash buffer 1, 2, 3, LiCl wash Buffer and TE buffer and eluted.

Following elution of histone mark and transcription factor samples, RNase (Roche cat# 10109169001) and Proteinase K (Invitrogen cat# 25530-049) treatments were performed, and reverse crosslinked at 65 °C overnight. DNA was purified with QiaQuick PCR Purification Kit (Qiagen, cat# 28106). Libraries were then prepared using the NEBNext Ultra II DNA library Prep Kit for Illumina (NEB cat# E7645S) with single (NEB cat# E7335, cat# E7710) or dual (NEB cat# E7600, cat# E7780) indexing. Two replicates were performed for each condition. Indexed libraries were validated for quality and size distribution using a TapeStation 4150 (Agilent), quantified by KAPA Library Quantification Kit (Roche cat# KK4824) and paired-end sequenced on Illumina NextSeq 550 (38bp+38bp) or Novaseq 6000 (61bp+61bp).

### In situ Hi-C

*In situ* Hi-C was performed in Granta519-Cas9 control and LRmCherry2.1-EBF1-g7 cells sorted 3 days post transduction. Briefly, 1x10^6^ cells were washed with 1X PBS containing 3% BSA and fixed with 2% formaldehyde and proceeded with Arima-HiC Kit (ARIMA cat# A510008) per manufacturer’s instructions. Samples passing Arima-QC1 were used for library construction with Accel-NGS 2S Plus DNA Library Kit (Swift cat# 21024) and 2S SET A INDEXING KIT (Swift cat# 26148) and assessed with Arima-QC2. Samples passing Arima-QC2 were subsequently PCR amplified based on Arima-QC2 calculations, quantified with TapeStation 4150 (Agilent) and KAPA Library Quantification Kit (Roche cat# KK4824) and paired-end sequenced on Illumina NextSeq 550 (38bp+38bp).

### SMC1 and EBF1 HiChIP

SMC1 and EBF1 HiChIP was performed in Granta519 as previously described with slight modifications^27^. Briefly, 2x10^7^ cells were crosslinked with 2% formaldehyde (Thermo Fisher Scientific, cat# 28908) for 10 min and subsequently quenched with 0.125 M glycine (Invitrogen, cat# 15527-013). Chromatin was digested using MboI restriction enzyme (NEB, cat# R0147), followed by biotin incorporation with Biotin-14-dATP (Jena Bioscience cat# NU-835-BIO14-S). DNA was then ligated with T4 ligase (NEB cat# M0202L) and sonicated on Covaris LE220. Sheared chromatin was 4-fold diluted with ChIP dilution buffer (16.7 mM Tris pH 7.5, 167 mM NaCl, 1.2 mM EDTA, 0.01% SDS, 1.1% Triton X-100), cleared with IgG (CST cat# 2729S) and then incubated with anti-SMC1 antibody (Bethyl, cat# A300-055A), polyclonal anti-EBF1 (1C) antibody recognizing an N-terminal EBF1 peptide (RG)^38^, or commercial EBF1 antibody (Millipore cat# AB10523) at 4 °C overnight. Chromatin-antibody complexes were captured by Protein-A magnetic beads (Pierce, cat# 88846) and subsequently washed with Low Salt Wash Buffer, High Salt Wash Buffer, LiCl Wash Buffer (detailed procedure available upon request) and eluted. DNA was purified with MinElute PCR Purification Kit (Qiagen, cat# 28004) and quantified using Qubit dsDNA HS Assay Kit (Invitrogen, cat# Q32851). 50-150ng DNA was used for capture with Dynabeads MyOne Streptavidin C-1 (Invitrogen, cat# 65001) and an appropriate amount of Tn5 enzyme (Illumina, cat# FC-121-1030) was added to captured DNA to generate sequencing library. Paired-end sequencing was performed on Illumina Novaseq 6000 (61bp + 61bp).

### Micro-C

Micro-C was performed as previously described with optimization^22^. For each reaction, 5x10^6^ cells were crosslinked with 1% formaldehyde for 10 min and quenched with 125 mM glycine for 5 min. Cells were washed with 1X PBS twice at room temperature and resuspended at 1x10^6^ cells/mL and crosslinked at a final concentration of 3 mM EGS (Thermo Scientific cat# PI21565). Double-crosslinking was quenched with 0.4 M glycine for 10 min at room temperature, washed twice with cold PBS, and stored at -80°C until further processing.

Thawed cell pellet was resuspended in 500 μL MB#1 buffer (50 mM NaCl, 10 mM Tris-HCl pH 7.5, 5 mM MgCl_2_, 1 mM CaCl_2_, 0.2 % NP-40 and 1X cOmplete EDTA-free PI) and rotated for 20 min at 4 °C. Nuclei were centrifuged at 8,000 G for 5 min at 4 °C and rinsed with 500 μL MB#1 once and resuspended in 500 μL MB#1. Chromatin is digested by appropriate amounts of MNase (25 U per 10^6^ for Granta519 and 20 U per 10^6^ for DND41) for 10 min at 37 °C shaking at 850 rpm. MNase digestion was quenched by 4 mM EGTA for 10 min shaking at 65 °C. Nuclei pellet was rinsed twice in 500 μL of cold MB#2 (50 mM NaCl, 10 mM Tris-HCl pH 7.5, 10 mM MgCl_2_), and 5 μL was saved and adjusted to 200 μL TE buffer and saved in -80 °C as 1% input. Digested chromatin was then subjected to de-phosphorylation with Shrimp Alkaline Phosphatase rSAP (NEB cat# M0371S), end-chewing with T4 Polynucleotide Kinase (NEB cat# M0201L), and end-labeling with Biotin-14-dATP (Jena Bio cat# NU-835-BIO14-L), and Biotin-11-dCTP (Jena Bio cat# NU-809-BIOX-L) (detailed procedure available upon request). Enzymatic reactions were quenched with 30 mM EDTA for 20 min at 65 °C and rinsed once with 500 μL of cold MB#3 (50 mM Tris-HCl pH 7.5, 10 mM MgCl_2_). Chromatin was then ligated with T4 DNA ligase at 20 U/μL for 4 hours with rotation at room temperature. After biotin-end removal with Exonuclease III for 15 min at 37°C, 25 μL 20 mg/mL Proteinase K and 25 μL 10% SDS were added to sample or input and de-crosslinked at 65 °C 850 rpm overnight.

After RNase A treatment at 37 °C for 45 minutes, chromatin was mixed with 1X volume of PCI (Invitrogen cat# 15593031) at room temperature and extracted with MaXtract high density columns (Qiagen cat# 129056). 0.1X volume of 3 M NaAc, 2.5X volume of 100% ethanol and 0.5 μL 20 mg/ml glycogen were added and precipitated at -80 °C for 2 hours. Chromatin was pelleted and resuspended with 50 µL 1X TE and then concentrated with ZymoClean DNA purification kit (cat# D4013). For each input and sample, 1 µL purified chromatin was examined with Agilent D5000 reagent and ScreenTape (cat# 5067-5588 and 5067-5589) and the remaining samples were separated in 3.5% NuSieve GTG Agarose gel (Lonza cat# 50080). Successful MNase digestion and T4 ligation should yield ∼70% mono-nuclei at 150 bp and di-nuclei at 300 bp on TapeStation, respectively. Ligated chromatin between 300 bp – 400 bp was excised from the gel and purified with Zymoclean Gel DNA Recovery Kit (cat# D4007) and eluted in 50 μL elution buffer. Samples were quantified with Qubit dsDNA HS Assay Kit and saved in -80 °C until further processing.

For each sample, 5 μL Dynabeads MyOne Streptavidin C1 beads were transferred to a fresh tube with 500 μL 1X TBW (5 mM Tris-HCl pH 7.5, 0.5 mM EDTA, 1 M NaCl, 0.05% Tween-20) and placed on magnetic rack. Beads were washed again with 1X TBW and resuspended in 150 μL 2X B&W buffer (10 mM Tris-HCl pH 7.5, 1 mM EDTA, 2 M NaCl). 100 μL elution buffer was added to each thawed chromatin sample and joined with beads and rotated at room temperature for 20 min. Beads were cleared on a magnet and resuspended in 300 μL 1X TBW shaking at 55°C for 900 rpm, 2 min and cleared on magnet. Beads were washed again with 1X TBW and 10 mM Tris-HCl and finally resuspended in 50 μL 0.1X TE. Beads containing chromatin were subject to end-prep and adaptor ligation following NEBNext Ultra™ II DNA Library Prep Kit (NEB cat# E7645S) and washed twice with 1 X TBW and 10 mM Tris-HCl before resuspension in 24 μL elution buffer. Minimum PCR with 2X KAPA HiFi Hot Start Mix (Roche cat# KK2601) and NEB dual indexes were performed with DNA on beads and purified twice with SPRIselect beads (Beckman Coulter cat# B23318), quantified with HSD1000 TapeStation 4150 (Agilent) and KAPA Library Quantification Kit (Roche cat# KK4824) and paired-end sequenced on Illumina Novaseq 6000 (61bp + 61bp).

### Optical Reconstruction of Chromatin Architecture (ORCA)

#### Probe Design and Synthesis

3 Mb genomic sequences spanning *MYC* promoter in hg19 (chr8: 127,800,000 – 130,800,000) were divided into 100 x 30 Kb bins/steps and extracted from Ensembl GRCh37.75. For each bin, multiple sets of 42 bp probes with homology to the genomic sequences were designed with OligoMiner pipeline^92^ using the default parameters except adjusting the spacing between probes. The set with closest to 200 probes was further manually filtered by NCBI BLAST to remove probes with potential multiple hybridization sites on the human genome. This ensures that the signal of each step of the sequential imaging is specific, and the intensity is comparable. For probes in each step, a unique 20 bp barcode sequence was added to both sides^31,93^. For all probes, a pair of 20 bp universal primer binding sequences were added for PCR amplification, resulting in final probe length of 122 bp. A total of 19,649 probes were designed for 3 Mb *MYC* locus and used in DND41 (**Fig. 7**). To further increase specificity, a subset of 5,891 probes covering 5’ of the *MYC* promoter were ordered as a separate pool and used in Granta519 (**Figs. 3 and 4**). A set of 8,128 probes and a set of 5,912 probes were designed for *KLF7* and *ZEB2* loci using similar procedures, respectively, and all probe sequences can be found in **Table S5**. All oligo pools were purchased from Twist.

Oligo pools were amplified based on Twist recommended PCR amplification protocol to achieve maximal yield without overamplification. For instance, 2 ng *MYC* 3 Mb probes were first amplified using universal primers with fiducial binding sequence (20 bp) added to both forward and reverse primers. KAPA HiFi HotStart ReadyMix (Roche cat# KM2605) was used with annealing temperature at 65 °C for 12 cycles per manufacturer’s recommended PCR procedure. PCR products were purified with Zymo DNA Clean & Concentrator-5 (cat# D4013) and quantified with Nanodrop. 1 μL purified DNA was ran on TapeStation HSD1000 to determine the uniform length of PCR products. Overamplification will lead to non-specific peaks larger than expected product (162 bp). Then 20 ng DNA per reaction was used for a second amplification with 6-10 reactions and examined again for uniform PCR product. Similar quality control procedure was done for *MYC* 5’ probes, *KLF7* probes and *ZEB2* probes. PCR products were joined and purified with Zymo DNA Clean & Concentrator-5 (cat #4013), and T7 in vitro transcription (NEB cat# E2040S) was performed at 37 °C overnight. Reverse transcription was performed with maxima RT-H (Thermo cat# EP0751) at 50 °C with universal forward primer for 2.5 hours, and EDTA/NaOH were added to remove RNA at 92 °C for 10 min. Finally, ssDNA was purified with Zymo DNA Clean & Concentrator-100 (cat# D4030), SpeedVac dried and resuspended in molecular grade water to 50 μM.

#### Primary Probe hybridization

40 mm round glass coverslips (Bioptechs cat# 10200-064) were coated with 0.01% Poly-L-Lysine (Sigma-Aldrich cat# P4832) for 15 min at room temperature. SSCT/F buffer containing 2X SSC + 0.1% Tween-20 + 50% formamide (20X SSC Buffer Corning cat# 46-020-CM, Tween-20 Millipore cat# 655204, deionized formamide Thermo cat# AM9344) were prepared at room temperature and preheated in 60 °C water bath and 95 °C water bath. A square region was drawn on each coverslip with a hydrophobic barrier pen (Vector Laboratories cat# H-4000), and 200 μL of Granta519 or DND41 cells at 10^7^ cells/mL in PBS were gently settled within the barrier. After 30 min incubation in a humidified chamber at room temperature, cells on coverslips were crosslinked with 4% formaldehyde in PBS (Thermo cat# 28908) and permeabilized for 15 min with 0.5% Triton-100 (Millipore cat# 648466) in PBS at room temperature. Cells were then dehydrated with 70%, 90% and 100% ethanol for 2 min each and slightly air dried before washing in SSCT/F for 5 min at RT, 3 min at 95 °C and 20 min at 60 °C. Primary hybridization mix containing 7 μL of 50 μM probes, 10 μL of 100 mM dNTPS (Denville cat# C788T68), 4 μL 10 mg/mL RNase A (Invitrogen cat# EN0531), 45 μL 100% formamide and 24 μL 4X dextran sulfate mix (40% dextran sulfate Sigma cat# D8906, 4% PVSA Sigma cat# 278424, 8X SSC, 0.4% Tween-20) were thoroughly mixed and added to each coverslip and sealed with a 22 x 30 mm coverslip. Coverslips were then heated on a heat block in 95 °C water bath for 3 min to denature genomic DNA and incubated overnight at 37 °C.

#### Sequential Imaging

To accurately select fields of view (FOV) and Z-stacks, hybridization of fiducial oligos and the first readout oligos was conducted before mounting the coverslip on the microscope, which differs from the original ORCA protocol. Specifically, the 22 x 30 mm coverslip was removed after overnight hybridization and the 40 mm round coverslip containing cells was washed in 2X SSCT at 60 °C and room temperature for 10 min each. For *MYC* 5’ probes in Granta519 and *ZEB2* probes in DND41, secondary hybridization mix containing 1 μL 100 μM Cy3-fiducial oligos, 1 μL 20 μM Readout_008 oligos, 2 μL 10 μM NDB_1281_Alexa647 oligos, 22.5 μL 100% formamide, 22.5 μL 4X dextran sulfate mix and 41 μL H_2_O were mixed and added per coverslip. For *MYC* 3 Mb probes in DND41, Readout_029 and NDB_1279_Alexa647 were used. For *KLF7* probes in DND41, Readout_005 and NDB_1279_Alexa647 were used. After 1h incubation in a humidified chamber at room temperature shielded from light, coverslips were washed in 2X SSCT at 60 °C and room temperature for 5 min each and transferred individually to 60mm Petri dishes containing 2X SSC. Coverslips were either used immediately or stored for less than 24 hours at 4 °C.

Sequential hybridization and imaging on the microscope (Bruker Vutara VXL, software version SRX 7.3.22) were proceeded after visual confirmation of even cell distribution, bright fiducial signal at 0.05% laser power and highly overlapping first readout signal at 0.5% laser power, which were the laser powers used for all ORCA experiments. After the first round of imaging, strand-displacement (also called toehold) oligos for the first readout were hybridized for 30 min in 25% formamide, 2X SSC and washed twice with 30% formamide in 2X SSC. Successful removal of signal was confirmed with imaging. The coverslip was then rinsed of imaging buffer using 2X SSC, and the second readout oligos were hybridized for 30 min. For each 30-step ORCA, we conducted 60 rounds of hybridization, washing, imaging for each set of readout and strand-displacement oligos, a process that took on average 1 week. Similarly, each 36-step and 72-step ORCA experiments were conducted with 72 with 144 rounds of sequential hybridization, washing, and imaging, and took on average 1 and 2 weeks, respectively. An automated fluidic system with daily refreshed buffers was used for all the experiments.

### Cell proliferation

Cell proliferation was measured with CellTiter Glo Luminescent Cell Viability Assay (Promega cat# G7571) according to the manufacturer’s instructions. Granta519-EBF1-FKBP-KI clone 27 was plated in 5 replicates with 1,000 cells per well on 96-well plates. Luminescence was measured 24 hours after plating (day 0) and every day for a total of 6 days. Statistics for cell growth changes were calculated using Student’s t-test.

### Flowcytometry analysis

Granta519-Cas9, JVM-2-Cas9 and PGA-1-Cas9 cells with control or LRmCherry2.1-EBF1-g7 were sorted 3 days post transduction and cultured for 3 days (day 6). For each replicate, 5x10^5^ cells were washed in 1X PBS and resuspended in 100 μl 1X Annexin V binding buffer (BD cat# 556454). 5 μL FITC-Annexin V (BD cat# 556420) was added and incubated in the dark at room temperature for 15min. 400 μL of 1X Annexin V binding buffer containing 1:1000 TO-PRO3 (ThermoFisherScientific cat# R37170) was added and immediately proceeded to Flow cytometry analysis on BD LSR II. 1x10^6^ Granta519 EBF1-FKBP-KI clones 27 and 97 were treated with 125 nM dTAG for 0, 6 and 24 hours and stained with L/D Aqua (Invitrogen cat# L34957) for 15 min at room temperature. Cells were washed with 1X PBS and then stained with 5 μL APC human anti-PD-L1 (BioLegend cat# 329708) in 100 μL 1X PBS for 15 min on ice before analysis on BD LSR II. Experiments were repeated three times with 3-5 replicates per condition.

### Western blot

0.2-1x10^6^ cells were washed in ice cold 1X PBS and lysed with whole lysis buffer (2% SDS, 60 mM Tris) supplemented with proteinase inhibitors. Protein concentration was determined with DC Protein Assay Reagents Package (Bio-rad cat# 5000116) and 5-20 μg proteins were used for SDS-PAGE electrophoresis in tris-mes-sds running buffer (GenScript cat# M00654, cat# M00677). Gels were then transferred to methanol activated PVDF membrane (Cytive cat# 10600023), blocked with 5% skim milk in 1X TBST at room temperature for 1 hour, and incubated with primary antibodies at 1:1,000 to 1:20,000 dilutions in 5% skim milk at 4 °C overnight. After washing with 1X TBST 3X 10min, secondary antibodies (Millipore cat# 401315-2ml, cat# 401215- 2ml) were added at 1:3,000 or 1:6,000 dilution in 5% skim milk and incubated at room temperature for 1 hour. Following 3 x 15min wash in 1X TBST, imaging was done with ECL Prime Western Blotting Detection Reagents (Cytiva cat# RPN2232) and autoradiography films. Primary antibodies: GAPDH (D16H11) XP (CST cat# 5174); MYC [Y69] (Abcam cat# ab32072); β-actin clone AC-74 (Sigma-Aldrich cat# A5316); Cas9 (7A9-3A3) (CST cat# 14697); SMC1 (BL-205- 2G8) (Bethyl cat# A700-018).

## QUANTIFICATION AND STATISTICAL ANALYSIS

The statistical significance of differences between measurements was determined by Wilcoxon rank sum test using R (version 3.6.1) wilcox.test function, unless otherwise stated. Statistical details of experiments can be found in figure legends. Visualizations were done with R.

### Definition of regulatory elements

The following definitions of regulatory elements were used throughout the manuscript. Promoters: promoters were defined as ± 2.5 kilobases (Kb) from the transcription start site (TSS) of each expressed gene. Enhancers: enhancers were defined as H3K27ac peaks excluding the ones overlapping with promoters.

### Gene annotation

A total of 2,828,317 Ensembl transcripts in GRCh37.75 assembly were downloaded in gtf format. For each Ensembl gene id (ENSG), the longest transcript (ENST) was used to assign a unique transcriptional start site and gene position. After exclusion of genes annotated as rRNA or on chromosome M, 57,209 gene annotations were used in RNA-seq analysis.

### ATAC-seq data analysis

#### Alignment

Reads from ATAC-seq experiments were trimmed with Trim Galore (version 0.4.1) with parameters -q 15 --phred33 --gzip --stringency 5 -e 0.1 --length 20. Trimmed reads were aligned to the Ensembl GRCh37.75 primary assembly including chromosomes 1-22, chrX, chrY, chrM and contigs using BWA (version 0.7.13)^94^ with parameters bwa aln -q 5 -l 32 -k 2 -t 6. Paired-end reads were grouped with bwa sampe -P -o 1000000. Reads mapped to contigs, ENCODE blacklist and marked as duplicates by Picard (version 2.1.0) were discarded and the remaining reads were used in downstream analyses and visualization.

#### Peak calling and Differential accessibility analysis

Peaks in each aligned replicate were identified using MACS2 (version 2.0.9)^95^ with parameters --nomodel --nolambda --format=BAM -g hs -p 1E-5 --bw=300 --keep-dup=1. All peaks from replicates of each experiment were combined using bedtools ‘merge’ function and the union of peaks was quantified over each aligned bam file using bedtools ‘coverage’ and normalized to RPKM. Differential accessibility analysis was performed using DEseq2^96^ with parameters test = “Wald”, betaPrior = F, fitType = “parametric”. For comparison of time-course EBF1 degradation and restoration, significance cutoff was abs(Log2(fold change)) > 1 and FDR < 1E-3.

### ChIP-seq data analysis

Reads from ChIP-seq experiments were aligned with the same procedure as ATAC-seq above. For Granta519 EBF1 ChIPseq, peaks of each library were identified using MACS2 (version 2.0.9)^95^- with parameters -p 1E-3 -g hs --nomodel --shiftsize=0.5*fragment_length --format=BAM--bw=300 --keep-dup=1 and with corresponding input control. 25,227 reproducible EBF1 ChIP-seq peaks identified in libraries of both replicates of both antibodies were used in downstream analyses. For comparing protein loading, reproducible EBF1-bound regions were quantified on each ChIP-seq bam files using bedtools ‘coverage’ and normalized to RPKM and averaged between the two replicates of each antibody. Clustering of reproducible EBF1-bound regions (**Fig. 1A**) was performed with R function hclust(dc, method=“average”, members=NULL) and hc_tree<- cutree(hc, k = 15) and visualized with ‘pheatmap’. Dynamic EBF1 peaks were filtered as reproducible EBF1 peaks > 1 averaged RPKM in untreated Granta519-EBF1-FKBP-KI clone 27. 7,777 peaks with Log2(fold change) < -1 of averaged RPKM between 6h and 24h dTAG-treated versus untreated and Log2(fold change) > 1 of averaged RPKM between 6h and 24h washout versus 24h dTAG-treated were defined as dynamic EBF1 peaks. SMC1, CTCF and YY1 ChIP-seq libraries were individually quantified on EBF1 peaks using bedtools ‘coverage’ and normalized to RPKM and averaged between the two replicates per condition, and Log2 fold changes were calculated as above.

For tag density plots, aligned bam files of the two replicates of each condition were merged using samtools (version 1.3)^97^ ‘cat’ command. For each merged library, fragment length was estimated with HOMER ‘makeTagDirectory’. HOMER ‘annotatePeaks.pl’ was used on merged libraries and visualized with R function ‘pheatmap’. For merged libraries genome tracks, bedgraph of reads normalized to reads per million (RPM) were generated with bedtools ‘genomecov’. Selected genomic loci were visualized with R package Sushi (version 1.18.0)^98^ function ‘plotBedgraph’. Genome-wide uploadable bigWig files were generated with UCSC tools (version 329)^99^ ‘bedGraphToBigWig’.

### Hi-C, SMC1 HiChIP and Micro-C analysis

#### Alignment and significant interaction calling

Raw reads for Granta519-Cas9 Ctrl and EBF1 KO Hi-C, Granta519-EBF1-FKBP-KI clones 27 and 97 time-course MicroC, and DND41 parental and GSI-resistant Micro-C samples were processed with Juicer (version 1.5.6)^100^ with parameters -g hg19 -s none. To generate contact maps of 30 Kb resolution in comparison with ORCA data, juicer_tools.1.7.6 ‘pre’ command was used on merged_nodups.txt with -r 2500000, 1000000, 500000, 250000, 100000, 50000, 30000, 25000, 10000, 5000 to generate the final .hic files. Normalized contact matrices were extracted using juicer_tools.1.7.6 ‘dump’ command with ‘observed VC_SQRT BP 30000’ for chromosome 8 (*MYC*) and chromosome 2 (*ZEB2* and *KLF7*) for plotting contact maps with R ‘pheatmap’ (**Figs. 3, 6 and 7**). Each .hic file was also converted to .mcool files using ‘hic2cool convert’ to maintain VC_SQRT normalization for quantification in **Fig. 2**.

Raw reads for each HiChIP sample were processed with HiC-Pro (version v2.5.0)^101^ to obtain putative interactions with default parameters except LIGATION_SITE = GATCGATC and GENOME_FRAGMENT generated for MboI restriction enzyme. Valid pairs (VI), self-circle (SC) and dangling-end (DE) interactions in cis were used as input for significant interaction calling in ‘.bedpe’ format. Mango (version 1.2.0)^102^ step 4 identified putative significant interaction anchors by MACS peak calling with MACS_qvalue = 0.05 and MACS_shiftsize = 75. Mango step 5 identified significant interactions with default parameters except maxinteractingdist = 2000000 and MHT = found. Only significant interactions with PETs >= 5 were used in the following analyses.

#### TAD boundary analysis

Raw reads for Granta519-Cas9 ctrl and EBF1 KO Hi-C samples were processed with open2c distiller-nf (version v0.3.4)^103^ to generate mcool files. Boundaries were identified in Python (version 3.9.13) using cooltools^104^ with window sizes of 50000,100000 and filtered with boundary strength > 0.5 with both window sizes. Boundaries in control conditions with EBF1 < 2.5 Kb are considered EBF1-bound. Pileup of EBF1-bound or EBF1-unbound boundaries in ctrl and EBF1 KO HiC were calculated with cooltools.pileup and visualized with matplotlib.pyplot (**Fig. 1E**).

#### Long-range regulatory loops

Long-range regulatory loops were defined using SMC1 and EBF1 HiChIP significant interactions. Significant interactions with EBF1, SMC1, CTCF, YY1 or H3K27ac ChIP-seq signal on both loop anchors were further classified into enhancer-enhancer (EE), enhancer-promoter (EP) promoter-promoter (PP), CTCF-CTCF and other interactions based on the presence of enhancers, promoters and CTCF peaks at the summit of the two anchors +/- 5 Kb. Normalized interactions at selected loci were visualized with R package Sushi^98^ (version 1.18.0) function ‘plotBedpe’.

#### Differential compartment calling

A/B compartments detected at 50 Kb resolution was tested. Hi-C processed data was converted into the HOMER (v4.11)^19^ ‘HiCSummary’ format. Subsequently, the Hi-C data was processed with HOMER command ‘makeTagDirectories’ with parameters -format ‘HiCSummary’. To identify compartment regions across the genome, HOMER runHiCpca.pl command was used on each condition with H3K27ac ChIP-seq as reference to ensure sign consistency via the -active option. To concatenate PC values from each condition, HOMER annotatePeaks.pl was used with the - noblanks and -bedGraph options. Scatterplot of the PC values from EBF1 Ctrl and KO conditions were plotted.

#### Hub analysis

Hubs were defined as previously described as “3D clique”^27^. Briefly, an undirected graph of regulatory interactions was constructed for each HiChIP where each vertex was an enhancer, or a promoter and each edge was a significant enhancer-enhancer, enhancer-promoter, or promoter-promoter interaction. Clusters of interacting enhancers and/or promoters were defined by spectral clustering^105^ of the regulatory graph interactions using cluster_louvain function in igraph R package with default parameters. Connectivity of an enhancer-promoter cluster was defined as the number of edges connecting vertices within the cluster. The connectivity of clusters was ranked in ascending order and plotted against the rank. Enhancer-promoter hubs were defined as clusters of enhancers and promoters with connectivity greater than the cutoff set as the elbow of the cumulative connectivity curve, which is shown as a tangent line in **Fig. 1F**. Hubs with > 10 or <= 10 expressed genes (defined in the following section) are considered gene-dense or gene-sparse hubs, respectively. Pairwise interactions between enhancers and promoters in hubs were quantified in Granta519-EBF1-FKBP-KI clone 27 time-course Micro-C .mcool files with VC_SQRT normalization using cooltools^104^. Interactions with Log2(fold change) < -0.5 between 6h or 24h dTAG-treated versus untreated and Log2(fold change) > 0.5 between 6h or 24h washout versus 24h dTAG-treated were defined as differential loops. Pileup plots of differential loops on Granta519-EBF1-FKBP-KI clone 27 time-course Micro-C were generated with R package GENOVA (version 1.0.1)^106^ APA and ‘visualise’ functions (**Fig. 2G**).

### RNA-seq data analysis

RNA-seq data was aligned to Ensembl GRCh37.75 primary assembly including chromosomes 1- 22, chrX, chrY, chrM and contigs using STAR (version 2.5)^107^ with parameters -- outFilterIntronMotifs RemoveNoncanonicalUnannotated --alignIntronMax 100000 -- outSAMstrandField intronMotif --outSAMunmapped Within --chimSegmentMin 25--chimJunctionOverhangMin 25. Strand-specific read counts were quantified using Subread (version 1.5.1) ‘featureCounts’ with parameters -t exon -g gene_id -s 1 -T 6 and used as input to differential gene expression analysis. Read counts were normalized to reads per million per kilobase (RPKM) for each gene. For Granta519-Cas9 ctrl and EBF1 KO samples, expressed genes were determined as genes with > 1 RPKM in at least 3 of the individual libraries. For Granta519-EBF1-FKBP-KI clones 27 and 97, expressed genes were determined as genes with > 1 average RPKM of the triplicates per condition in at least 3 conditions per clone. Pairwise differential gene expression analysis was performed using DEseq2^96^ with parameters test = “Wald”, betaPrior = F, fitType = “parametric”. Significance cutoff was abs(Log2(fold change)) > 0.5 and FDR < 0.05. Expressed genes in gene-dense and gene-sparse hubs were ranked by averaged RPKM and the top 500 genes of each category were used to conduct Gene Ontology (GO) enrichment analysis. GO sets only enriched in gene-dense or gene-sparse hubs with FDR < 1E-7 were visualized in R (**Figs. 1K and 1L**).

### ORCA data analysis

#### ChrTracer3, QC and imputation

Raw images were exported as individual .tiff per FOV, step, z-plane and channel using Vutara SRX (No hardware version 7.1.38). Tiff files for each readout were compiled as ConvZscan.dax and ConvZscan.inf files using MATLAB as input for ChrTracer3^31^. ChrTracer3 was followed stepwise for drift correction, spot selection, and spot fitting using default parameters except changing the pixel to nm conversion for xy and z planes. Traces identified in all FOV per experiment were concatenated and further analyzed in R and MATLAB. For each trace, the number of registered steps and brightness (column “h” in ChrTracer3 output table) were used for quality control. For 30-step walks, traces with less or equal to 5 missing steps were kept. For 36- step walks, the cutoff was 6 missing steps and for 72-step walks the cutoff was 15.

#### Distance and contact frequency

Granta519-EBF1-FKBP-KI *MYC* locus: pairwise distances were calculated on unimputed traces per condition (untreated, 6h dTAG, 24h dTAG, 6h washout, 24h washout) per biological replicate (clone 27 and clone 97). Using the median distance of all neighboring steps (i.e. step 1 to 2, 2 to 3, … 29 to 30) as cutoff, the contact frequency vectors of unimputed traces were generated and the Pearson correlations were calculated. After confirmation of >95% correlation between the two replicates, the traces were combined for downstream analyses. More details can be found in **Table S6.** The median distances across combined traces excluding missing values were used to plot the final distance map per condition (**Fig. S3**). The neighboring step cutoff was recalculated with combined traces and used to generate contact frequency maps (**Fig. 3**) and compare enhancer-promoter contact frequencies (**Fig. 4**). Interaction between *MYC* and super-enhancer E1 was defined as distances between either step 25-7 or 25-8 smaller than cutoffs per condition. *MYC* to super-enhancer E2 interaction was defined as distance between step 25-11 smaller than cutoffs per condition. For E1, equally spaced bins were [1-18, 2-19, …, 8-25, …,13-30] or [2-18 (for padding), 1-19, 2-20, …, 7-25, …, 12-30] (**Fig. 4B**). For E2, equally spaced bins were [4-18, 5-19, …, 11-25, …, 16-30] (**Fig. 4C**). Control regions were defined as [6-23, 7-24, 9-26] or [7-23, 8-24, 10-26] for *MYC*-to-E1 and [10-23, 11-24, 13-26] for *MYC*-to-E2 and the contact frequencies were averaged with missing values removed (**Figs. 4D and 4E**). Error bars in **Fig. 4** were 95% confidence intervals of contact frequencies calculated from 1,000 sets of 500 randomly sampled alleles.

DND41 *KLF7* and *ZEB2* loci: pairwise distances were calculated on unimputed traces per condition (GSI-sensitive, GSI-resistant). The median distances across combined traces excluding missing values were used to plot the final distance map per condition (**Figs. 6 and S6**). 250 nm was used as cutoff for determining pairwise interaction. DND41 *MYC* locus: pairwise distances were calculated on unimputed traces per condition (GSI-sensitive, GSI-resistant) in two biological replicates and combined. The median distances across combined traces excluding missing values were used to plot the final distance map per condition (**Fig. S7**). The contact frequency matrices were generated using the median distance of all neighboring steps per condition as cutoffs per condition. *MYC* and super-enhancer TE1 was defined as interacting whenever any distance between step 4 to steps 52, 53, or 54 was smaller than the condition’s cutoff. *MYC* and super-enhancer TE2 was defined as interacting whenever any distance between step 4 to steps 65, 66, or 67 was smaller than the condition’s cutoff. Error bars in **Fig. 7** were 95% confidence intervals of contact frequencies calculated from 1,000 sets of 1,000 randomly sampled alleles.

#### Clustering and radius of gyration

Granta519 EBF1-FKBP-KI *MYC* locus: Filtered, imputed alleles from all conditions and replicates were combined and the pairwise distance vectors were calculated as input to K-means clustering using R ‘kmeans’ function with centers = 10 and iter.max = 20. The neighboring step cutoff was recalculated with combined traces and used to generate contact matrices for each K. Contact maps of each K were manually inspected and arranged as in **Fig. 3F**. K-means clustering was then conducted on each condition and manually inspected to be arranged in the same order as **Fig. 3F**. Stratification of whether the allele has E1, E2 and *MYC* interactions were determined as described above using the unimputed values of the allele. Error bars in **Figs 4H and S4E-H** were confidence intervals calculated from 1000 sets of 500 randomly sampled alleles. Imputed alleles from all conditions were used for radius of gyration calculation per definition in **Fig. 4F**.

### Polymer Simulations

Simulation of loop extrusion was performed using Open2C’s polychrom Python library (v0.1.1)^108^ with slight modifications. The simulation framework is divided into two steps: a 1-dimensional step and a 3-dimensional step. The 1-dimensional step tracks the movement of 𝑐 × 2 cohesin legs/anchors (𝑐 = the number of cohesins) across a polymer comprised of 𝑁 monomers, where each monomer represents a genomic region of length 𝑀. The length in base pairs represented by the polymer can then be calculated as 𝐿 = 𝑁 × 𝑀. A predefined number of cohesins 𝑐 can each be either randomly loaded onto any monomer with a uniform probability distribution (‘random loading’), or loaded onto a specific region / sequence of monomers with uniform probability from which it will extrude (‘targeted loading’), these behaviors constitute the modified portion of the polychrom package. Targeted loading was primarily used in the *loading scenario* to simulate EBF1’s loading of cohesin at specific locations, while random loading was utilized in the *blocking scenario*. At each 1D step, the two anchors of the cohesin can take 4 possible actions: (1) unimpeded cohesin movement, which is defined as translocation of the anchors at positions 𝑥_1_ and 𝑥_2_ to their adjacent monomers 𝑥_1_ − 1 and 𝑥_2_ + 1 followed by forming a synthetic chemical bond between monomors occupied by the anchors, (2) unload the entire cohesin, including both anchors, from the polymer with probability [inline] where 𝑙 is the average number of monomers traversed (‘lifetime’) and load another cohesin in the same manner, (3) become blocked by a blocking element (i.e. CTCF or EBF1 as a blocker) at a monomer with probability 𝑝*_r_*, which may then release the anchor with probability 𝑝*_b_* at subsequent steps, or (4) become stalled against another cohesin anchor, in which case extrusion is also halted, but the lifetime 𝑙 of the cohesin is reduced by one-tenth (i.e. 𝑙*_stalled_* = 0.1 × 𝑙). The 3-dimensional step reads the cohesin positions produced by the 1-dimensional step and maps them onto a polymer in 3D space created with OpenMM. The polymer was considered as a self-avoiding walk (SAW) grown on a cubic lattice, and undergoes energy minimization followed by subsequent steps of cohesin movement as defined above. All thermodynamic and integration parameters were kept as the defaults provided by the polychrom library. The full list of altered simulation parameters distinguishing *loading* from *blocking scenarios* can be found in **Table S7**. The outputs of the 3-dimensional steps are the (x,y,z) coordinates of each monomer in the ensemble of simulated polymers, which are used in subsequent analyses to calculate distances and interaction frequencies in **Fig. 5**.

