## Supplementary Data for "Lineage-determining transcription factors constrain cohesin to drive multi-enhancer oncogene regulation"

**Affiliations:**
1. Department of Pathology and Laboratory Medicine,
2. Department of Genetics,
3. Department of Bioengineering,
4. Institute for Immunology and Immune Health
5. Epigenetics Institute, University of Pennsylvania, Philadelphia, PA, USA
6. Department of Biochemistry and Molecular Biology, University of Texas Medical Branch, Galveston, TX, USA
7. Max Planck Institute of Immunobiology and Epigenetics, Freiburg, Germany.

* Equal contributions
### co-corresponding authors

### Figure S1

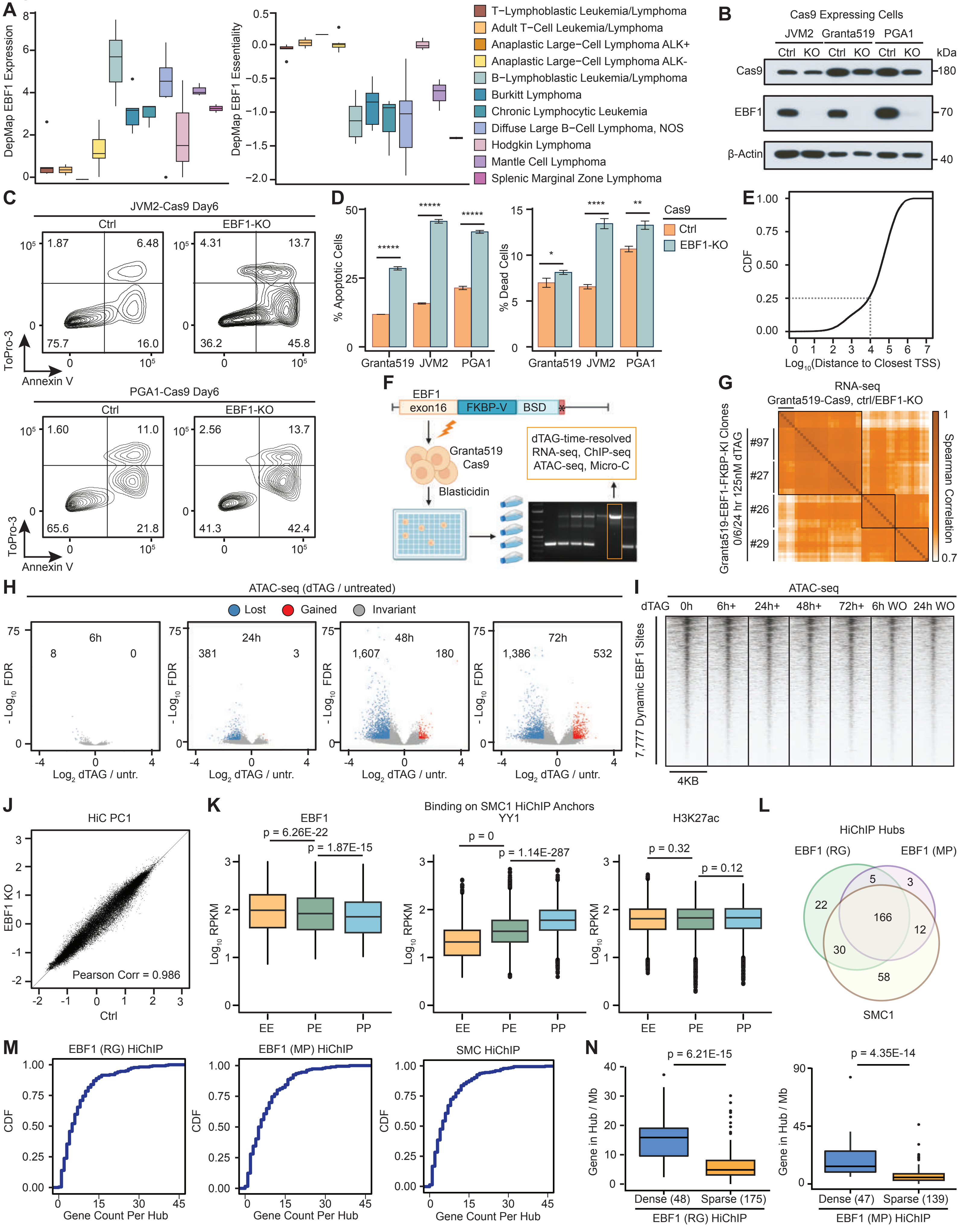

A
DepMap EBF1 Expression
DepMap EBF1 Essentiality
T−Lymphoblastic Leukemia/Lymphoma
Adult T−Cell Leukemia/Lymphoma
Anaplastic Large−Cell Lymphoma ALK+
Anaplastic Large−Cell Lymphoma ALK-
B−Lymphoblastic Leukemia/Lymphoma
Burkitt Lymphoma
Chronic Lymphocytic Leukemia
Diffuse Large B−Cell Lymphoma, NOS
Hodgkin Lymphoma
Mantle Cell Lymphoma
Splenic Marginal Zone Lymphoma
B
Cas9 Expressing Cells
JVM2 Granta519 PGA1
Ctrl KO Ctrl KO Ctrl KO
kDa
Cas9 180
EBF1 70
β-Actin 40
C
JVM2-Cas9 Day6
Ctrl
EBF1-KO
1.87 6.48 75.7 16.0
4.31 13.7 36.2 45.8
ToPro-3
Annexin V
PGA1-Cas9 Day6
Ctrl
EBF1-KO
1.60 11.0 65.6 21.8
2.56 13.7 41.3 42.4
D
% Apoptotic Cells
% Dead Cells
Granta519 JVM2 PGA1
Cas9
Ctrl
EBF1-KO
E
CDF
Log10(Distance to Closest TSS)
F
EBF1
exon16
FKBP-V
BSD
Granta519
Cas9
Blasticidin
dTAG-time-resolved
RNA-seq, ChIP-seq
ATAC-seq, Micro-C
G
RNA-seq
Granta519-Cas9, ctrl/EBF1-KO
Granta519-EBF1-FKBP-KI Clones
0/6/24 hr 125nM dTAG
#97
#27
#26
#29
Spearman Correlation
H
ATAC-seq (dTAG / untreated)
Lost
Gained
Invariant
6h 8 0
24h 381 3
48h 1,607 180
72h 1,386 532
- Log10 FDR
Log2 dTAG / untr.
I
ATAC-seq
dTAG 0h 6h+ 24h+ 48h+ 72h+ 6h WO 24h WO
7,777 Dynamic EBF1 Sites
4KB
J
HiC PC1
EBF1 KO
Ctrl
Pearson Corr = 0.986
K
Binding on SMC1 HiChIP Anchors
EBF1
p = 6.26E-22
p = 1.87E-15
YY1
p = 0
p = 1.14E-287
H3K27ac
p = 0.32
p = 0.12
Log10 RPKM
EE PE PP
L
HiChIP Hubs
EBF1 (RG)
EBF1 (MP)
SMC1
22 5 3 166 12 30 58
M
EBF1 (RG) HiChIP
EBF1 (MP) HiChIP
SMC HiChIP
CDF
Gene Count Per Hub
N
p = 6.21E-15
p = 4.35E-14
Gene in Hub / Mb
Dense (48) Sparse (175)
EBF1 (RG) HiChIP
Dense (47) Sparse (139)
EBF1 (MP) HiChIP

**Figure S1: EBF1 is required for MCL growth but not for maintenance of chromatin accessibility or compartments.**

A: Boxplots depicting higher expression (left) and essentiality (right) of EBF1 in Non-Hodgkin B lymphomas compared to T-cell leukemia/lymphoma from DepMap.

B: Western blotting of EBF1 in Cas9-expressing JVM2, Granta519 and PGA1 MCL showing efficient depletion of EBF1 three days after transfection of sgRNA targeting EBF1. β-actin is loading control.

C, D: Flow cytometry plots (C) and quantification (D) of cell apoptosis and death measured by Annexin V and ToPro-3 staining. Ctrl and EBF1-KO Cas9-expressing MCL cells were sorted three days post lentiviral transduction and cultured for three days. Representative experiment of 3 biological replicates with 5 technical replicates. Student's t-test P-value: *p < 0.05, **p < 0.01, ****p < 0.0001, *****p < 0.00001.

E: Cumulative distribution plot (CDF) of distances between each reproducible EBF1 peak to the closest expressed gene TSS in Granta519 showing > 75% EBF1 peaks are located 10 Kb away from TSS.

F: Schematics of knocking in $FKBP^{F36V}$ domain at the stop codon of endogenous EBF1 gene in Granta519-Cas9 cells. Single cell clones were selected with blasticidin and successful insertion of the $FKBP^{F36V}$ cassette was validated with genomic DNA PCR. Clones with efficient EBF1 degradation after $dTAG^{V}$-1 treatment (referred to as dTAG) are used.

G: Hierarchical clustering of normalized reads (RPKM) from Ctrl and EBF1-KO Granta519-Cas9, and EBF1-FKBP-KI clones 26, 27, 29, 97 with 0, 6, 24-hour 125 nM dTAG treatment showing higher similarity of clones 27 and 97 with Granta519-Cas9 transcriptome.

H: Volcano plot showing ATAC-seq signal fold enrichment (x axis) versus false discovery rate (FDR) (y axis) in Granta519 EBF1-FKBP-KI clone 27 treated with 125 nM dTAG for 6, 24, 48, 72 hours compared with untreated cells. Each point depicts an accessible element, color coded by blue, red, and black based on significantly decreased, increased, or unchanged accessibility in treated cells, respectively. Significance cutoff: FDR < 1E-5 and Log2(fold change) => 1.

I: Heatmaps displaying overall unchanged normalized accessibility levels at dynamic EBF1 binding sites. Each column of ATAC-seq signals is centered on bona fide EBF1-bound elements per Figure 1D +/- 2 Kb flanking sequences with 50 bp resolution.

J: Scatterplot of PC1 values of Hi-C contact matrices in Ctrl and EBF1-KO Granta519-Cas9 cells showing largely invariable A/B compartments.

K: Boxplots of EBF1 (left), YY1 (middle) and H3K27ac (right) loading at the promoters (P) and enhancers (E) interacting with significant long-range DNA loops of SMC1 HiChIP in Granta519.

EBF1 preferentially binds enhancers, while YY1 preferentially binds promoters. H3K27ac is negative control. P-values: Wilcoxon rank sum test.

L: Venn diagram comparing overlapping of the genomic coordinates of enhancer-promoter hubs in Granta519 SMC1 and EBF1 (RG and MP) HiChIP showing high concordance of hubs identified by three assays.

M: Cumulative distribution plots showing expressed genes count in each enhancer-promoter hub defined by EBF1 (RG and MP) and SMC1 HiChIP.

N: Boxplots showing higher number of genes per megabase in gene-dense compared to gene-sparse hubs defined by EBF1 HiChIP using RG (left) and MP (right) antibodies. P-values: Wilcoxon rank sum test.

### Figure S2

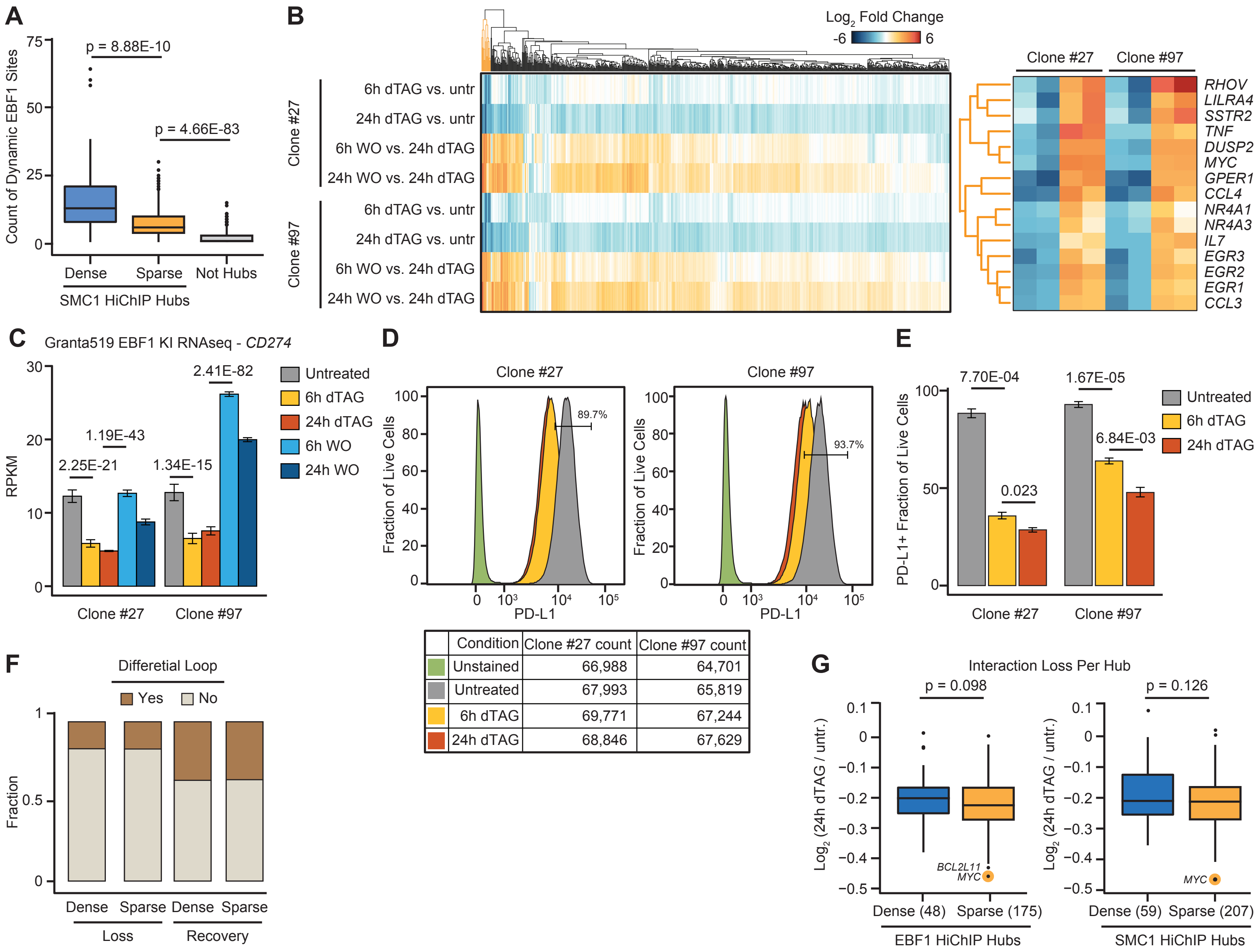


| Condition | Clone #27 count | Clone #97 count |
|---|---|---|
| Unstained | 66,988 | 64,701 |
| Untreated | 67,993 | 65,819 |
| 6h dTAG | 69,771 | 67,244 |
| 24h dTAG | 68,846 | 67,629 |

**Figure S2: EBF1 rapidly alters expression of *CD274*, a gene located in a gene-sparse hub.**

A: Boxplots comparing number of EBF1 binding sites in gene-dense, gene-sparse and non-hub regions, as determined with SMC1 HiChIP. P-values: Wilcoxon rank sum test.

B: Left: Hierarchical clustering of Log2(fold change) between indicated conditions showing high concordance of differentially expressed genes in clones 27 and 97. Right: Zoomed in of the gene cluster highlighted in yellow in the left panel.

C: Barplots of normalized RNA-seq reads showing significant *CD274* (encoding PD-L1) down- and up-regulation 6 and 24 hours after dTAG treatment and washout, respectively, in Granta519 EBF1-FKBP-KI clones 27 and 97. Data represent mean ± S.D. of 3 replicates per condition. FDR from DESeq2.

D, E: Flow cytometry plots (D) and quantification (E) of PD-L1-positive live cells in clones 27 and 97 showing down-regulation of PD-L1 after EBF1 degradation. Lower panel of (D) indicates total live cell counts per condition by L/D aqua staining showing cell viability is unaffected by dTAG treatment in 24 hours. Upper panel of (D) shows a representative of 3 biological replications.

F: Barplots showing the proportion of enhancer-promoter Micro-C interactions with more than 2-fold interaction frequency changes in gene-dense and gene-sparse hubs after dTAG treatment and washout.

G: Boxplots showing significantly stronger loss of Micro-C interactions among enhancers and promoters 24 hours after dTAG treatment in gene-sparse compared to gene-dense hubs as defined by EBF1 (left) and SMC1 (right) HiChIP. Each dot represents the average Log2(fold change) of 24h washout over 24h dTAG of interactions in each hub.

#### Figure S3

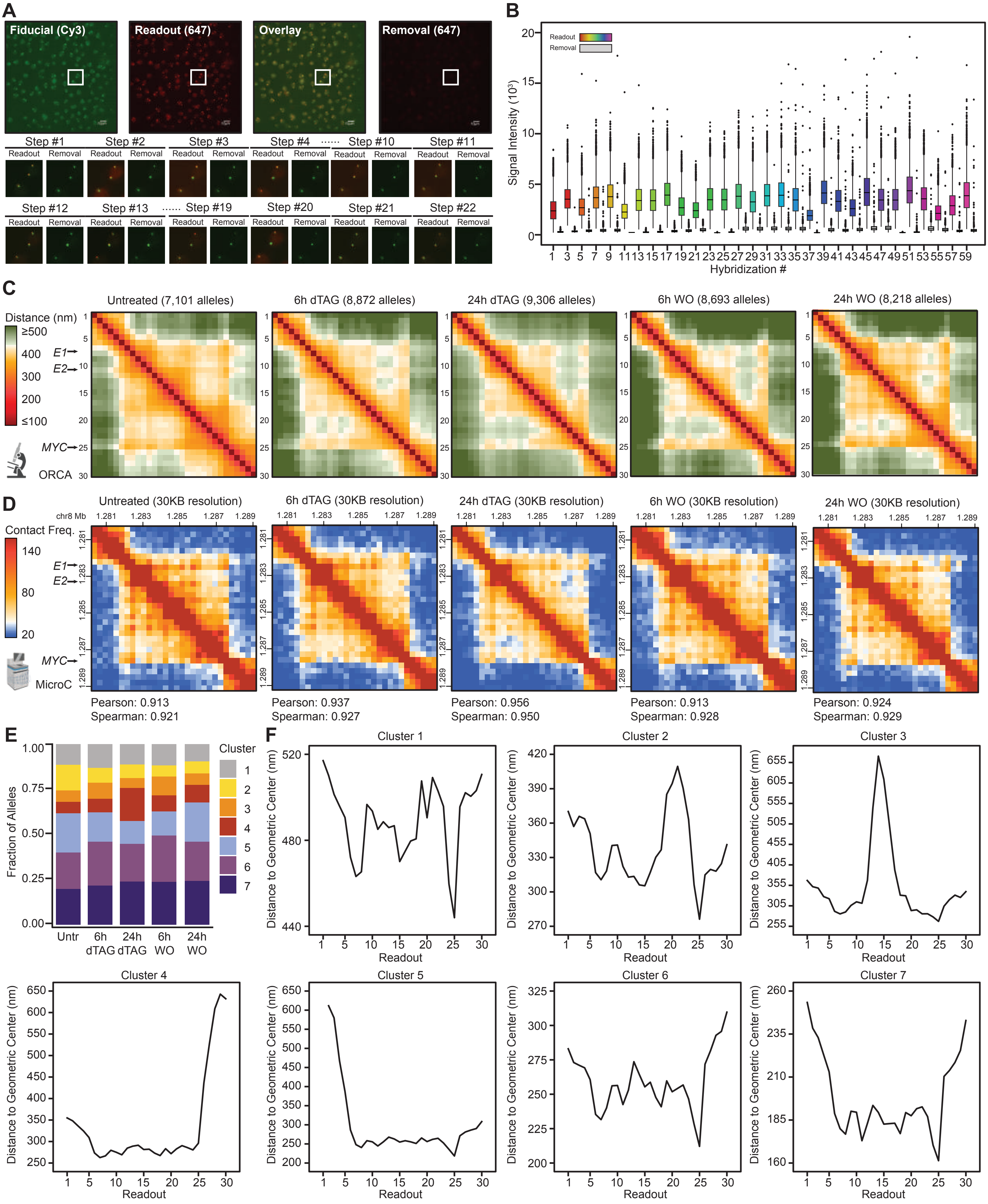
A
Fiducial (Cy3)
Readout (647)
Overlay
Removal (647)
Step #1
Step #2
Step #3
Step #4
Step #10
Step #11
Step #12
Step #13
Step #19
Step #20
Step #21
Step #22
Readout
Removal
B
Readout
Removal
Signal Intensity (10³)
Hybridization #
C
Untreated (7,101 alleles)
6h dTAG (8,872 alleles)
24h dTAG (9,306 alleles)
6h WO (8,693 alleles)
24h WO (8,218 alleles)
Distance (nm)
≥500
400
300
200
≤100
E1
E2
MYC
ORCA
D
Untreated (30KB resolution)
6h dTAG (30KB resolution)
24h dTAG (30KB resolution)
6h WO (30KB resolution)
24h WO (30KB resolution)
chr8 Mb
Contact Freq.
140
80
20
MicroC
Pearson: 0.913
Spearman: 0.921
Pearson: 0.937
Spearman: 0.927
Pearson: 0.956
Spearman: 0.950
Pearson: 0.913
Spearman: 0.928
Pearson: 0.924
Spearman: 0.929
E
Fraction of Alleles
Cluster
Untr
6h dTAG
24h dTAG
6h WO
24h WO
F
Cluster 1
Cluster 2
Cluster 3
Cluster 4
Cluster 5
Cluster 6
Cluster 7
Distance to Geometric Center (nm)
Readout

**Figure S3: High quality ORCA chromatin traces show EBF1-dependency of common topological conformations at the *MYC* locus.**

A: Top: example images of nuclei on a field of view of the hybridization and subsequent removal of the first readout. Bottom: zoomed-in images show the process of hybridization and removal at the indicated steps for example alleles showing consistent signals throughout the experiment.

B: Boxplots of signal intensities (column h in ChrTracer 3 output) at each hybridization and removal indicating strong signal-to-noise ratio.

C: Time-course ORCA pairwise distance maps showing gradual increase and decrease of pairwise distances at the *MYC* locus after dTAG treatment and washout. Each point represents the median of pairwise distances between two steps across all alleles. Each condition combines the alleles from clones 27 and 97. Alleles are not imputed, and missing values are excluded from the calculations.

D: Time-course Micro-C interaction frequency maps of clone 27 at the *MYC* locus at 30 Kb resolution to match with the step size of ORCA. Pearson and Spearman correlations of each condition are calculated between vectorized Micro-C and ORCA maps in Figure 3D.

E: Barplots showing the fraction of alleles in each of the *MYC* locus common topological conformations per condition.

F: Distances of each segment to the geometric centers of the traces. The median across all alleles per common topological conformations in Figure 3F are calculated and plotted for each segment of the *MYC* ORCA experiment.

#### Figure S4

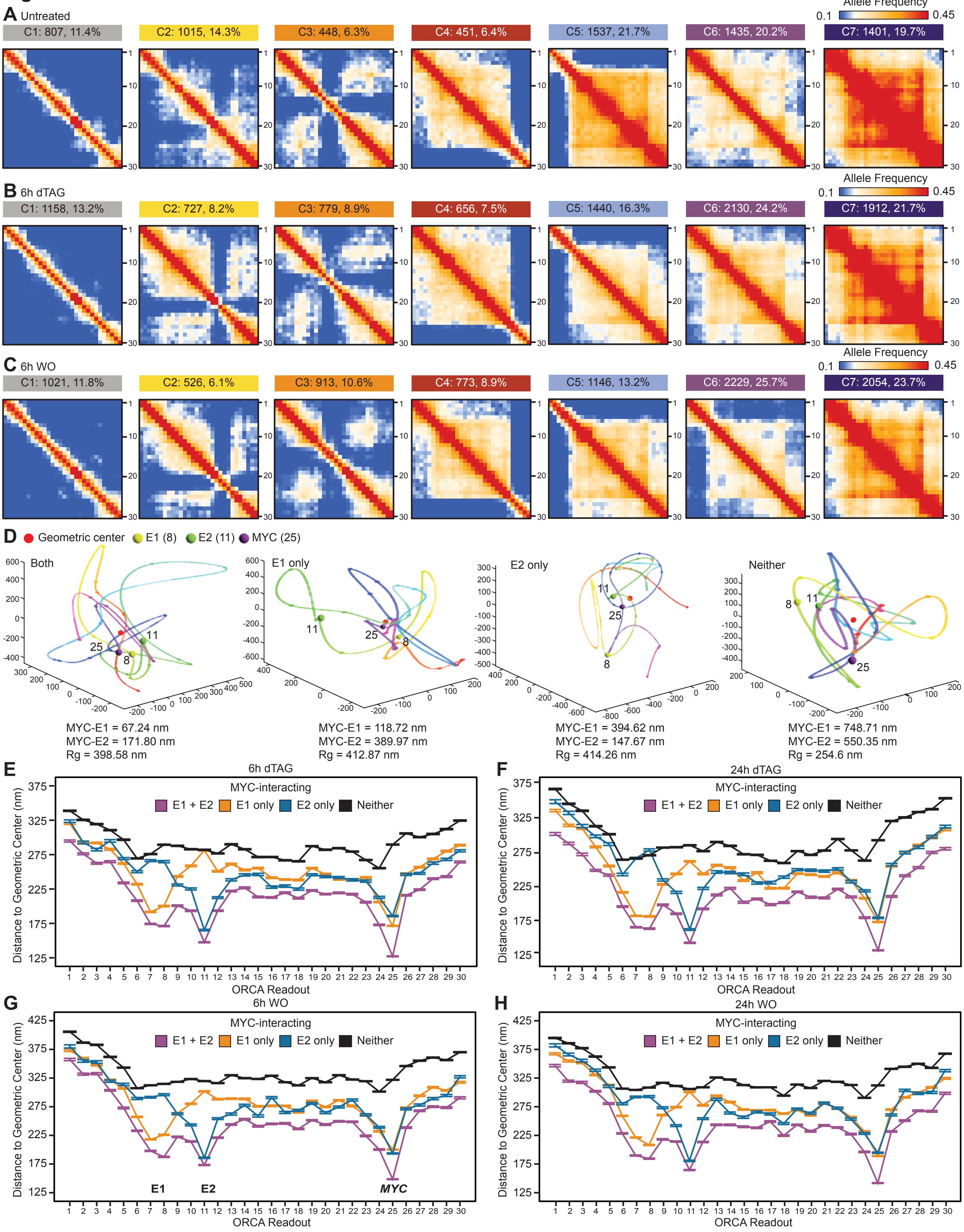
A Untreated
Allele Frequency
0.1
0.45
C1: 807, 11.4%
C2: 1015, 14.3%
C3: 448, 6.3%
C4: 451, 6.4%
C5: 1537, 21.7%
C6: 1435, 20.2%
C7: 1401, 19.7%
B 6h dTAG
C1: 1158, 13.2%
C2: 727, 8.2%
C3: 779, 8.9%
C4: 656, 7.5%
C5: 1440, 16.3%
C6: 2130, 24.2%
C7: 1912, 21.7%
C 6h WO
C1: 1021, 11.8%
C2: 526, 6.1%
C3: 913, 10.6%
C4: 773, 8.9%
C5: 1146, 13.2%
C6: 2229, 25.7%
C7: 2054, 23.7%
D Geometric center E1 (8) E2 (11) MYC (25)
Both
MYC-E1 = 67.24 nm
MYC-E2 = 171.80 nm
Rg = 398.58 nm
E1 only
MYC-E1 = 118.72 nm
MYC-E2 = 389.97 nm
Rg = 412.87 nm
E2 only
MYC-E1 = 394.62 nm
MYC-E2 = 147.67 nm
Rg = 414.26 nm
Neither
MYC-E1 = 748.71 nm
MYC-E2 = 550.35 nm
Rg = 254.6 nm
E 6h dTAG
F 24h dTAG
G 6h WO
H 24h WO
MYC-interacting
E1 + E2
E1 only
E2 only
Neither
Distance to Geometric Center (nm)
ORCA Readout
E1
E2
MYC

**Figure S4: Time-course ORCA data in MCL show interaction of the *MYC* promoter and super enhancers at the allelic geometric centers.**

A-C: Allele frequency maps of traces for each of the 7 common topological conformations of the *MYC* locus in untreated (A), 6-hour dTAG treated (B) and 6-hour dTAG washout (C) showing reproducibility of the observed conformations. Clustering and calculation of frequencies are performed on pairwise distances matrices of imputed alleles.

D: Example reconstructed chromatin traces for alleles with *MYC*-E1-E2 three-way interaction, *MYC*-E1 interaction, *MYC*-E2 interaction and no interaction emphasizing the central positioning of interacting elements. Note that the allele with no *MYC*, E1, E2 interactions has the smallest radius of gyration among the examples.

E-H: Distances of each segment to the geometric centers of the traces with *MYC*-E1-E2, only *MYC*-E1, only *MYC*-E2 or no *MYC* promoter-enhancer interaction (Neither). The median values of stratified alleles are calculated and plotted for each segment of the ORCA experiment in 6-hour dTAG treated (E), 24-hour dTAG treated (F), 6-hour dTAG washout (G) and 24-hour dTAG washout (H) cells. Error bars show 95% confidence interval from bootstrapping.

### Figure S5

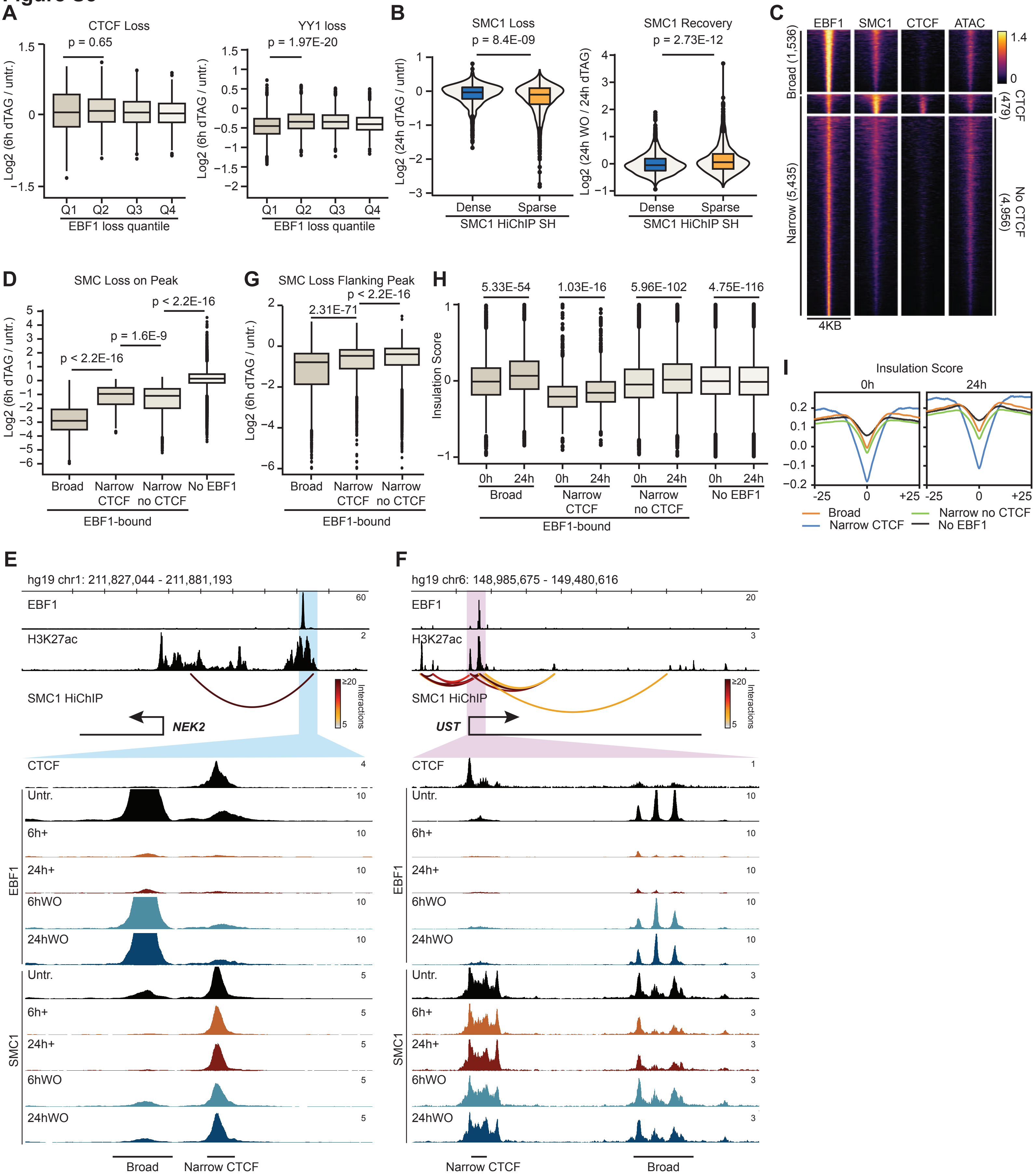

A
CTCF Loss
p = 0.65
Log2 (6h dTAG / untr.)
Q1 Q2 Q3 Q4
EBF1 loss quantile
YY1 loss
p = 1.97E-20
B
SMC1 Loss
p = 8.4E-09
Log2 (24h dTAG / untrl)
Dense Sparse
SMC1 HiChIP SH
SMC1 Recovery
p = 2.73E-12
Log2 (24h WO / 24h dTAG)
C
EBF1 SMC1 CTCF ATAC
Broad (1,536)
CTCF (479)
Narrow (5,435)
No CTCF (4,956)
4KB
D
SMC Loss on Peak
p < 2.2E-16
p = 1.6E-9
Log2 (6h dTAG / untr.)
Broad
Narrow CTCF
Narrow no CTCF
No EBF1
EBF1-bound
G
SMC Loss Flanking Peak
2.31E-71
p < 2.2E-16
H
5.33E-54
1.03E-16
5.96E-102
4.75E-116
Insulation Score
0h 24h
I
Insulation Score
0h
24h
-25 0 +25
Broad
Narrow CTCF
Narrow no CTCF
No EBF1
E
hg19 chr1: 211,827,044 - 211,881,193
EBF1
H3K27ac
SMC1 HiChIP
NEK2
≥20
Interactions
CTCF
Untr.
6h+
24h+
6hWO
24hWO
SMC1
Broad
Narrow CTCF
F
hg19 chr6: 148,985,675 - 149,480,616
UST

**Figure S5: EBF1 changes SMC1 but not CTCF and YY1 levels in MCL.**

A: Boxplots showing limited correlation between CTCF (left) and YY1 (right) ChIP-seq signals in 6h dTAG treatment (left) and washout (right) per quartile of EBF1 loading changes 6h after dTAG treatment.

B: Box and violin plots comparing differential SMC1 ChIP-seq signals in 24h dTAG treatment (left) and washout (right) at SMC1 peaks located in gene-sparse and gene-dense hubs. P-values: Wilcoxon rank sum test.

C: Heatmaps showing broad and focal EBF1 binding sites with a small subset co-occupied by CTCF. Clusters are generated and plotted with deepTools.

D: Boxplots showing stronger loss of SMC1 levels at broad EBF1-binding sites 6 hours after dTAG treatment. EBF1-binding sites as grouped per (C). P-values: Wilcoxon rank sum test.

E, F: Top three genomic tracks showing EBF1 and H3K27ac ChIP-seq as well as SMC1 HiChiP at *NEK2* (E) and *UST* (F) loci in untreated cells showing examples of broad and narrow EBF1 binding sites. Lower genomic tracks showing zoomed in EBF1, SMC1, and CTCF biding sites at *NEK2* (E) and *UST* (F) loci in the noted conditions demonstrating marked EBF1-dependency of SMC1 levels at the broad compared to narrow EBF1 binding sties, and no change at the CTCF sites.

G: Boxplots showing significant reduction in SMC1 levels at the broad EBF1 binding sites compared to narrow EBF1 binding sites with or without CTCF. EBF1 peaks +/- 1500 bp flanking sequences are considered.

H, I: Boxplots (H) and pileup plots (I) showing significant reduction in insulation potential at broad EBF1 binding sites 24 hours after dTAG treatment compared to no EBF1, narrow EBF1 with CTCF, and narrow EBF1 without CTCF sites.

### Figure S6

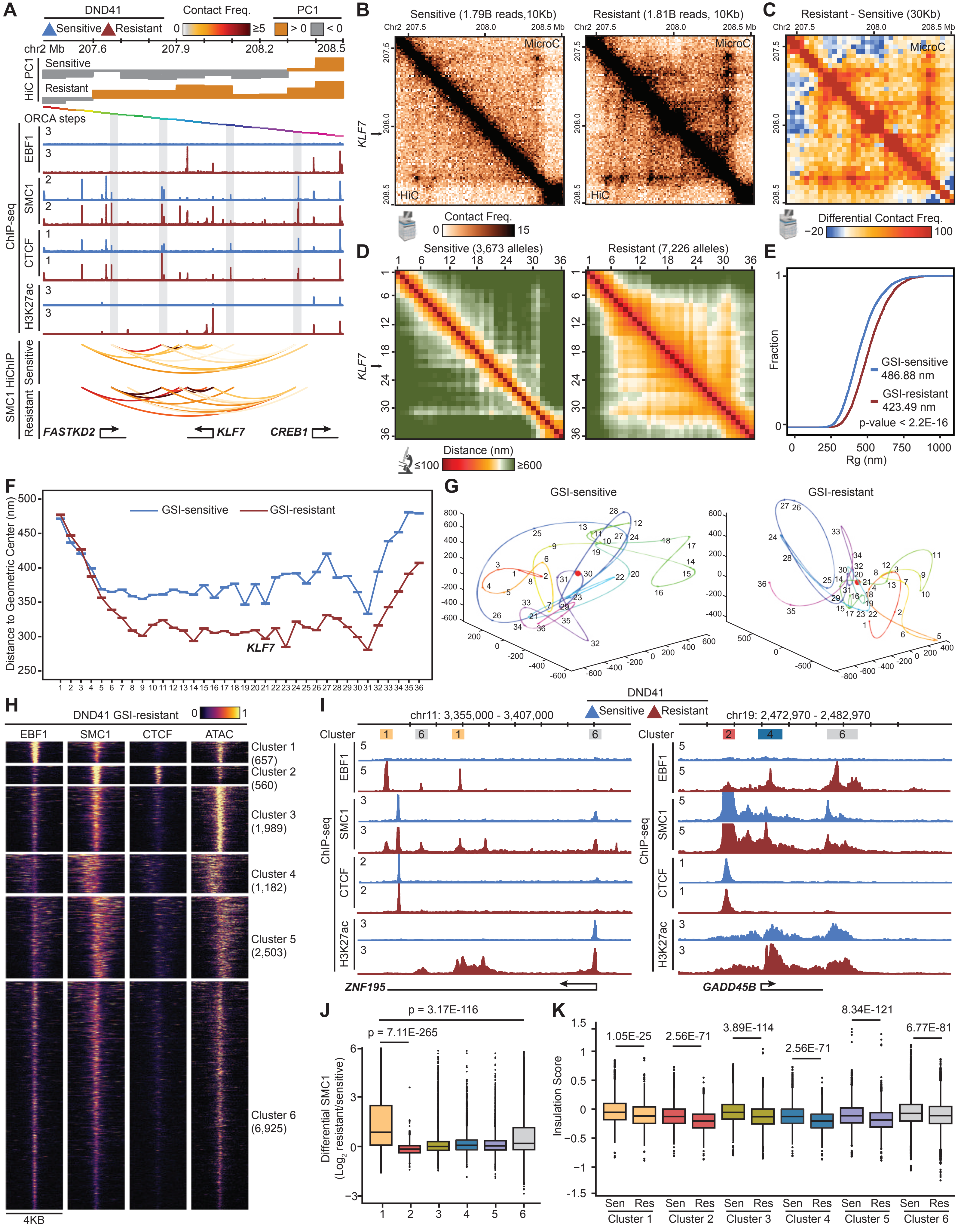

A
DND41
Sensitive
Resistant
Contact Freq.
0
≥5
PC1
> 0
< 0
chr2 Mb 207.6 207.9 208.2 208.5
HiC PC1
Sensitive
Resistant
ORCA steps
ChIP-seq
EBF1
SMC1
CTCF
H3K27ac
SMC1 HiChIP
FASTKD2
KLF7
CREB1
B
Sensitive (1.79B reads,10Kb)
Resistant (1.81B reads, 10Kb)
MicroC
HiC
Contact Freq.
0
15
C
Resistant - Sensitive (30Kb)
Differential Contact Freq.
−20
100
D
Sensitive (3,673 alleles)
Resistant (7,226 alleles)
KLF7
Distance (nm)
≤100
≥600
E
Fraction
GSI-sensitive
486.88 nm
GSI-resistant
423.49 nm
p-value < 2.2E-16
Rg (nm)
F
Distance to Geometric Center (nm)
GSI-sensitive
GSI-resistant
KLF7
G
GSI-sensitive
GSI-resistant
H
DND41 GSI-resistant
EBF1
SMC1
CTCF
ATAC
Cluster 1 (657)
Cluster 2 (560)
Cluster 3 (1,989)
Cluster 4 (1,182)
Cluster 5 (2,503)
Cluster 6 (6,925)
4KB
I
DND41
Sensitive
Resistant
chr11: 3,355,000 - 3,407,000
chr19: 2,472,970 - 2,482,970
Cluster
ChIP-seq
EBF1
SMC1
CTCF
H3K27ac
ZNF195
GADD45B
J
p = 3.17E-116
p = 7.11E-265
Differential SMC1
(Log2 resistant/sensitive)
K
Insulation Score
1.05E-25
2.56E-71
3.89E-114
2.56E-71
8.34E-121
6.77E-81
Sen
Res
Cluster 1
Cluster 2
Cluster 3
Cluster 4
Cluster 5
Cluster 6

**Figure S6: EBF1 binds and activates *KLF7* promoter without changing its local radial positioning in GSI-resistant T-ALL.**

A: ChIP-seq tracks showing gain of active histone marks H3K27ac, EBF1, CTCF and SMC1 at *KLF7* locus in GSI-resistant DND41. Normalized SMC1 HiChIP arches showing gain of interactions in GSI-resistant DND41. The PC1 values of Hi-C contact correlation matrices in GSI-sensitive and GSI-resistant cells showing shift from B (<0) to A (>0) compartment. Rainbow-colored bars indicate the 36 steps of ORCA experiments. Bottom track indicates the positions of expressed genes.

B: Normalized Micro-C (top) and Hi-C (bottom) interaction frequency maps showing gain of interactions at the *KLF7* locus in GSI-resistant (right) compared to GSI-sensitive (left) DND41.

C: Differential (GSI-resistant minus GSI-sensitive) normalized Micro-C interaction maps with 30 Kb resolution matching ORCA at the *KLF7* locus showing gain of stripes in GSI-resistant DND41.

D: ORCA pairwise distance maps of *KLF7* chromatin traces in GSI-sensitive (left) and GSI-resistant (right) DND41. Each point represents the median of pairwise distances between two segments across all alleles. Alleles are not imputed, and missing values are excluded from the calculation.

E: Cumulative distributions of radius of gyration of *KLF7* chromatin traces in GSI-sensitive and GSI-resistant DND41 showing overall compaction of the *KLF7* locus in GSI-resistant cells. P-value: Kolmogorov–Smirnov test.

F: Distances of each segment to the geometric centers of the chromatin traces of *KLF7* locus in GSI-sensitive and GSI-resistant DND41. The median values of chromatin traces are calculated and plotted for each ORCA segment in GSI-sensitive and GSI-resistant cells. Error bars show 95% confidence interval from bootstrapping. Note lack of preferential positioning of the *KLF7* promoter (step 21) at the geometric centers of alleles in GSI-resistance.

G: Example reconstructed chromatin traces for alleles in GSI-sensitive (left) and GSI-resistant (right) *KLF7* emphasizing lack of central positioning of the gene promoter.

H: Heatmaps showing broad and focal EBF1 binding sites in GSI-resistant DND41 with a small subset co-occupied by CTCF. Clusters are generated and plotted with deepTools.

I: Genome tracks of EBF1, SMC1, CTCF and H3K27ac ChIP-seq in GSI-sensitive (blue) and GSI-resistant (red) DND41 at *ZNF195* (left) and *GADD45B* (right) loci showing examples of EBF1-bound elements for clusters noted in (H).

J: Boxplot showing Log2(fold change) of GSI-resistant vs GSI-sensitive SMC1 loading at each of the six clusters of EBF1 binding sites in (H). P-values: Wilcoxon rank sum test.

K: Boxplots showing gain of insulation potentials in GSI-resistance at each of the six clusters of EBF1 binding sites per (H). P-values: Wilcoxon rank sum test.

#### Figure S7

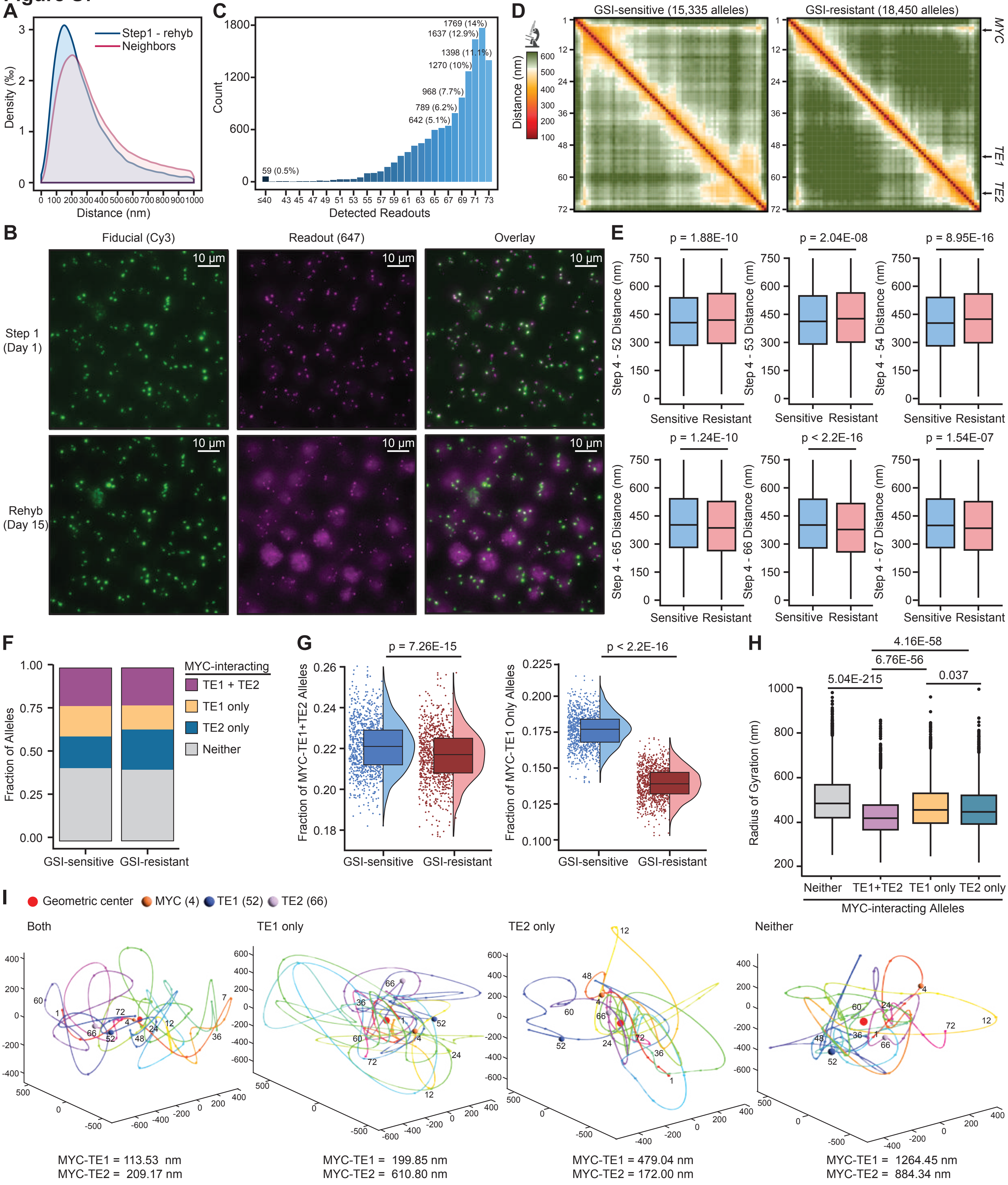

A
Step1 - rehyb
Neighbors
Density (‰)
Distance (nm)
C
Count
Detected Readouts
59 (0.5%)
642 (5.1%)
789 (6.2%)
968 (7.7%)
1270 (10%)
1398 (11.1%)
1637 (12.9%)
1769 (14%)
D
GSI-sensitive (15,335 alleles)
GSI-resistant (18,450 alleles)
Distance (nm)
MYC
TE1
TE2
B
Fiducial (Cy3)
Readout (647)
Overlay
Step 1 (Day 1)
Rehyb (Day 15)
10 µm
E
p = 1.88E-10
p = 2.04E-08
p = 8.95E-16
p = 1.24E-10
p < 2.2E-16
p = 1.54E-07
Step 4 - 52 Distance (nm)
Step 4 - 53 Distance (nm)
Step 4 - 54 Distance (nm)
Step 4 - 65 Distance (nm)
Step 4 - 66 Distance (nm)
Step 4 - 67 Distance (nm)
Sensitive Resistant
F
MYC-interacting
TE1 + TE2
TE1 only
TE2 only
Neither
Fraction of Alleles
GSI-sensitive GSI-resistant
G
p = 7.26E-15
p < 2.2E-16
Fraction of MYC-TE1+TE2 Alleles
Fraction of MYC-TE1 Only Alleles
GSI-sensitive GSI-resistant
H
4.16E-58
6.76E-56
5.04E-215
0.037
Radius of Gyration (nm)
Neither TE1+TE2 TE1 only TE2 only
MYC-interacting Alleles
I
Geometric center
MYC (4)
TE1 (52)
TE2 (66)
Both
TE1 only
TE2 only
Neither
MYC-TE1 = 113.53 nm
MYC-TE2 = 209.17 nm
MYC-TE1 = 199.85 nm
MYC-TE2 = 610.80 nm
MYC-TE1 = 479.04 nm
MYC-TE2 = 172.00 nm
MYC-TE1 = 1264.45 nm
MYC-TE2 = 884.34 nm

**Figure S7: High-quality 72-step tracing of the *MYC* locus in DND41 shows EBF1-dependency of *MYC* three-way interactions with its super-enhancers.**

A: Distributions of distances between all neighboring steps (red), and between step 1 and step 73 which is the rehybridization of readout 1 at the end of 15-day ORCA experiment in GSI-sensitive DND41 (blue) showing intact and immobile chromatin during the experiment.

B: Example field of view showing comparable signal levels in fiducial and readout channels at step 1 and 73 (i.e. rehybridization of readout 1).

C: Barplots showing distribution of allele counts with more than half of the readouts detected including rehybridization.

D: ORCA pairwise distance maps of *MYC* chromatin traces in GSI-sensitive (left) and GSI-resistant (right) DND41. Each point represents the median of pairwise distances between two segments across all alleles. Alleles are not imputed, and missing values are excluded from the calculation.

E: Boxplots showing increased pairwise distances between *MYC* (step 4) and TE1 (steps 52 – 54), and decreased pairwise distances between *MYC* (step 4) and TE2 (steps 65 – 67) in GSI-resistant DND41. P-values: Wilcoxon rank sum test.

F: Barplots showing fraction of alleles with *MYC* interacting with TE1, TE2, both TE1 and TE2 (TE1 + TE2) or neither in GSI-sensitive and GSI-resistant DND41.

G: Box and violin plots showing decrease in allele frequency with *MYC*-TE1-TE2 (left) and *MYC*-TE1 (right) interactions in GSI-resistant DND41. Each point represents the allele frequencies calculated from 1,000 randomly sampled alleles with a total of 1,000 rounds of random sampling per condition. P-values: Wilcoxon rank sum test.

H: Boxplots showing distributions of radius of gyration of alleles with *MYC*-TE1-TE2, only *MYC*-TE1, only *MYC*-TE2 or no *MYC* promoter-enhancer interaction (Neither) in GSI-sensitive DND41. P-values: Wilcoxon rank sum test.

I: Example reconstructed chromatin traces for alleles with *MYC*-TE1-TE2 three-way or only *MYC*-TE1 pairwise interactions from GSI-sensitive DND41, and only *MYC*-TE2 interaction or no promoter-enhancer interactions (Neither) from GSI-resistant DND41 emphasizing the central positioning of interacting elements.

**Supplemental Table Legends**

**Table S1:** Quantification of histone mark ChIP-seq signals at EBF1 peaks and coordinates of EBF1 dynamic peaks. Related to Figure 1.

**Table S2**: Differential gene expression in Granta519-Cas9, Granta519 EBF1-FKBP-KI clones 27 and 97. Related to Figures 1 and 2.

**Table S3**: Differential chromatin accessibility in Granta519 EBF1-FKBP-KI clone 27 during EBF1 degradation and recovery. Related to Figures 1.

**Table S4**: Hi-C A/B compartments, Hi-C TAD boundaries and HiChIP enhancer-promoter hubs in Granta519. Related to Figures 1 and 2.

**Table S5**: Probe sequences of all ORCA experiments. Related to Figures 3, 4, 6 and 7.

**Table S6**: Summary statistics of all ORCA experiments and traces from Granta519 EBF1-FKBP-KI untreated cells. Related to Figures 3, 4, 6 and 7.

**Table S7**: Parameters for polymer simulations. Related to Figure 5.
